# A Human Tumor-Immune Organoid Model of Glioblastoma

**DOI:** 10.1101/2025.06.16.660009

**Authors:** Shivani Baisiwala, Elisa Fazzari, Matthew X Li, Antoni Martija, Daria J Azizad, Lu Sun, Gilbert Herrera, Trinh Phan, Amber Monteleone, David A Nathanson, Anthony Wang, Won Kim, Richard G Everson, Kunal S Patel, Linda M Liau, Robert M Prins, Aparna Bhaduri

**Affiliations:** Department of Neurosurgery, UCLA Health, Los Angeles, CA, USA; Department of Biological Chemistry, David Geffen School of Medicine, University of California, Los Angeles, CA, USA; Department of Molecular and Medical Pharmacology, University of California, Los Angeles, CA, USA

## Abstract

A major obstacle to identifying effective therapies for the aggressive brain tumor glioblastoma is the lack of human-specific, immunocompetent models that reflect the human tumor microenvironment. To address this, we developed the immune-Human Organoid Tumor Transplantation (iHOTT) model. This is an autologous co-culture platform that integrates patient-derived tumor cells and matched peripheral blood mononuclear cells (PBMCs) within human cortical organoids, enabling the study of the patient-specific immune response to the tumor and tumor-immune interactions. This platform preserves tumor and immune populations, immune signaling, and cell-cell interactions observed in patient tumors. Treatment of iHOTT with pembrolizumab, a checkpoint inhibitor, mirrored cell type shifts and cell interactions observed in patients. TCR sequencing further revealed pembrolizumab-driven expansion of stem-like CD4-T-cell clonotypes exhibiting patient-specific repertoires. These findings establish iHOTT as a physiologically relevant platform for exploring autologous tumor–immune interactions and underscore the critical need for antigen-targeted strategies to enhance immunotherapy in glioblastoma.

## INTRODUCTION

Glioblastoma is the most common primary malignant adult brain tumor with a dismal prognosis for patients despite maximal standard of care therapy (1,2). Glioblastoma displays extraordinary cellular heterogeneity, but it remains unclear which populations drive therapeutic resistance or are clinically relevant. This knowledge gap is especially pronounced in the context of immunotherapy. Glioblastoma employs multiple immune evasion mechanisms - including expression of checkpoint signals, recruitment of immunosuppressive myeloid cells, and establishment of a dysfunctional immune microenvironment (3,4). While T cell-mediated approaches, including immune checkpoint inhibitors, DCVax, and CAR-T therapies, have shown significant promise in other cancers, there has been limited success with these strategies in glioblastoma (5). For example, only a small subset of patients show any objective response to checkpoint inhibition, and there is limited evidence as to what predicts response (6). Thus, models that faithfully preserve and permit manipulation of T cells as well as other immune cell types within the tumor context are urgently needed to accelerate therapeutic development (3).

Studying tumor-immune interactions requires a model system that supports both compartments. Most glioma models rely on orthotopic xenografts in immunodeficient mice, which are inherently limited by the need to use immunodeficient hosts (7–9). While genetically engineered mouse models allow immune studies, they differ substantially from human tumors in cell type composition, cytokine signaling, and overall tumor-immune microenvironment. Notably, mouse cytokines do not accurately mimic human cytokine interactions (10). Thus, a human-specific co-culture model that captures tumor-immune dynamics is essential for investigating resistance mechanisms and for identifying patient-specific responses to immunotherapy.

Recent work has incorporated CAR-T therapies into glioblastoma organoids, providing a platform for interrogating patient-specific responses to these T-cell-based therapies. While these systems provide valuable insights, they rely on organoids formed directly from tumor cells and immune populations already present within the tumor (11–14). Since glioblastoma is considered an immunologically “cold” tumor, these infiltrating immune cells are relatively limited (5). There remains an opportunity to study cellular interactions between tumor cells and a broader range of circulating immune cells in the context of the cortical brain environment. Furthermore, while existing 2D co-culture models and gliomasphere systems provide simpler platforms to study tumor-immune interactions, they inherently lack the cell-type diversity and complex multicellular interactions achieved by 3D organoid systems. 3D models offer critical advantages by capturing adaptations and resistance mechanisms that emerge from a more physiologically relevant and complex cellular environment.

We have developed a novel human organoid tumor transplantation (HOTT) model, which uses tumor cells directly derived from patient surgical resections, grown within stem cell-derived cortical organoids that simulate the human brain microenvironment (15). This system retains the cellular heterogeneity of the tumor and faithfully replicates key features of the tumor microenvironment (15). Building on this platform, here, we introduce immune cells into the system, creating the iHOTT (immune-HOTT) model. This innovation enables the study of complex interactions between autologous tumor cells and immune populations within a controlled human context. The flexibility of the iHOTT model allows for interrogation of specific immune and tumor cell subtypes, providing a unique opportunity to explore mechanisms of immune resistance and identify novel therapeutic targets.

In this study, we demonstrate how leveraging circulating PBMCs can be used to model a patient-specific immune response to tumors. Glioblastoma is a hypervascular tumor characterized by abnormal blood vessel networks, allowing continuous infiltration of circulating immune cells (16,17). Therefore, introducing freshly isolated and matched PBMCs provides a biologically relevant context for studying co-culture interactions. Unlike tumor-infiltrating immune cells, which may already be functionally exhausted, PBMCs represent a responsive circulating immune population encountering the tumor (18,19). This supports the utility of PBMC-based systems for modeling tumor-immune interactions and capturing relevant clonal dynamics in response to immunotherapy.

Evaluating the co-cultured cells reveals dynamic tumor-driven immune modulation and reflects how circulating immune cells respond upon entering the tumor microenvironment. We provide a detailed characterization showing that PBMC cell populations can be maintained in the iHOTT system, and that both cytokines and gene expression programs related to immune activation are generated specifically in the tumor-immune co-culture condition. By benchmarking the response of immune cells within iHOTT to that of actual patients treated with the checkpoint inhibitor pembrolizumab, we demonstrate that iHOTT recapitulates key cell types, cell-cell communication, and TCR clonotype features observed in patients. Collectively, our findings establish iHOTT as a powerful and versatile platform for studying human, patient-specific tumor-immune interactions and for the future identification of personalized immunotherapy approaches for glioblastoma.

## RESULTS

### iHOTT Enables Tumor-Immune Co-Culture

iHOTT enables the co-culture of freshly isolated, patient-matched glioblastoma cells and peripheral blood mononuclear cells (PBMCs). First, human cortical organoids are grown for 8-12 weeks as previously described [**Supplementary Figure 1A**] (20). These organoids contain major cell types present in the human cortex and thus provide a human cortical microenvironment for the subsequent transplant of patient tumor cells (20). On the day of surgery, the patient’s tumor is dissociated and infected with green fluorescent protein (GFP)-expressing lentivirus. These cells are transplanted at a 1:1 ratio with matched PBMCs onto our previously established cortical organoids [**Figure 1A**] (15). The media and culture conditions were optimized as described in *iHOTT Model Generation and Culture Methods* specifically to maximize recovery of immune subpopulations. Results showed that a 50-50 mixed media provided optimal immune subpopulation recovery despite a slightly reduced overall cell recovery **[Supplementary Figure 1B-C]**. Cells are cultured under these optimized media conditions for 7 days before harvest for sorting and single-cell RNA sequencing (scRNAseq), flow cytometry, and immunostaining. The media is additionally collected for cytokine profiling [**Figure 1A]**.

**Figure 1:**
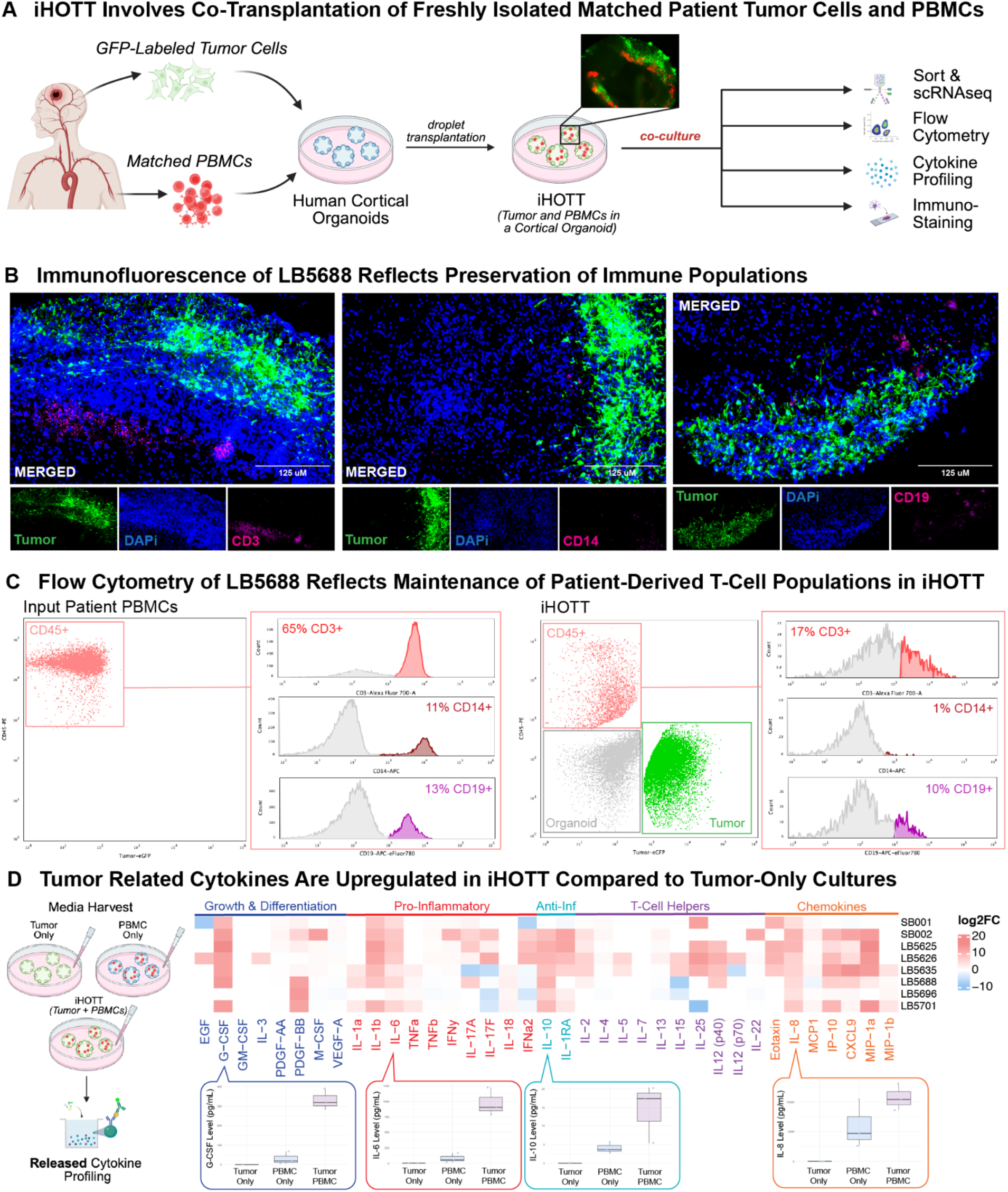
iHOTT Enables Tumor and Immune Co-Culture and Reflects Patient Tumor Cytokine Profiles A) Human cortical organoids derived from embryonic stem cells were cultured for 8–12 weeks before co-transplantation. Patient-derived glioblastoma cells were dissociated, transduced with GFP-expressing lentivirus, and co-transplanted with matched freshly isolated CellTrace-labeled patient PBMCs onto the organoid surface. Cultures were maintained for 7 days before processing. The inset shows representative fluorescence imaging of GFP+ tumor cells and CellTrace+ PBMCs within the organoid. After 7 days, organoids were collected for downstream flow cytometry, immunostaining, cytokine profiling, and single-cell RNA sequencing of tumor and immune compartments. B) iHOTT organoids were fixed, embedded in OCT, sectioned, and stained for GFP, CD3, CD14, and CD19. Immunofluorescence revealed robust infiltration of CD3+ T cells with minor populations of CD14+ myeloid and CD19+ B cells. GFP+ tumor cells were observed invading into the organoid. C) Flow cytometry of input PBMCs prior to culture revealed a predominantly CD45+ population composed primarily of CD3+ T cells. Analysis of dissociated iHOTT samples after 7 days showed maintenance of tumor (GFP+), organoid (GFP and CD45 negative), and immune (CD45+) compartments, with the immune population again composed primarily of CD3+ T cells, mirroring the original patient PBMC composition. D) Cytokine profiling of conditioned media from iHOTT, tumor-only, and PBMC-only cultures was performed at day 7. The heatmap shows log2 fold change in iHOTT compared to the tumor alone. iHOTT samples showed elevated secretion of tumor-associated cytokines, including G-CSF, IL-6, and IL-10, relative to both tumor-only and PBMC-only controls. Notably, PBMC-only cultures failed to produce these cytokines at comparable levels, indicating a tumor-immune interaction–dependent response within the iHOTT system.

Immunostaining of iHOTT was performed to evaluate the presence of tumor cells (eGFP), T cells (CD3), myeloid cells (CD14), and B cells (CD19) **[Figure 1B, Supplementary Figure 2A-B]**.

Complementary flow cytometry was conducted to quantify these populations within the CD45+ compartment using the same markers, and to evaluate the tumor cell engraftment [**Figure 1C]**. Results revealed an input PBMC composition of approximately 65% CD3+ T cells, 11% CD14+ myeloid cells, and 13% CD19+ B cells, consistent with previously reported distributions in human peripheral blood (21). Analysis of iHOTT confirmed tumor cell infiltration into the organoid. Furthermore, within the immune compartment, we observed recovery of CD3+ T cells and CD19+ B cells with a relatively smaller percentage of CD14+ myeloid cells **[Figure 1C]**. These percentages do not sum to 100%, likely due to the presence of other immune cell types not profiled with our marker panel.

### Cytokines Characteristic of the Tumor Microenvironment are Secreted from iHOTT

Media was collected from iHOTT as well as from tumor-only and PBMC-only organoid controls. Samples were analyzed for an extensive array of 37 cytokines and chemokines. Compared to tumor-only organoid cultures, iHOTT showed significant upregulation of cytokines previously associated with glioblastoma, including IL-6, IL-8, IL-10, and G-CSF **[Figure 1D].** IL-6 and IL-8 are pro-inflammatory cytokines, activating neutrophils and T cells. IL-10 is seen in inflammatory reactions, and G-CSF supports recruitment and survival of available myeloid cells in the microenvironment (22–25). Notably, for the majority of cytokines, increases in expression were not observed in the PBMC-only organoid condition, underscoring the critical role of tumor– immune cell interactions in driving cytokine production in this system [**Supplementary Figure 3A-B]**.

### scRNAseq Reflects Preservation of Tumor and Immune Cell Types

Three freshly-isolated patient tumors were co-cultured in iHOTT with matched PBMCs. Following co-culture, samples were sorted for eGFP⁺ (tumor cells) and CD45+ (immune cells), and each fraction was subjected to single-cell RNA sequencing as described in our *Methods*. Cell type annotations were performed using our previously established glioblastoma meta-atlas and an existing PBMC cell type atlas and were simplified as described in *Supplementary Table 1* (26,27). These annotations reflected preservation of all major tumor and immune cell types typically seen in patients (26,27) **[Figure 2A].** To support these assignments, canonical marker genes were visualized for tumor cells, specific immune subsets, immune activation, and exhaustion. Notably, immune cells in iHOTT showed limited expression of exhaustion markers such as HAVCR2 and LAG3, alongside robust expression of activation markers including GZMB and CD69, suggesting that iHOTT immune cells maintain an activated and functionally competent state rather than exhibiting signs of terminal exhaustion.

**Figure 2:**
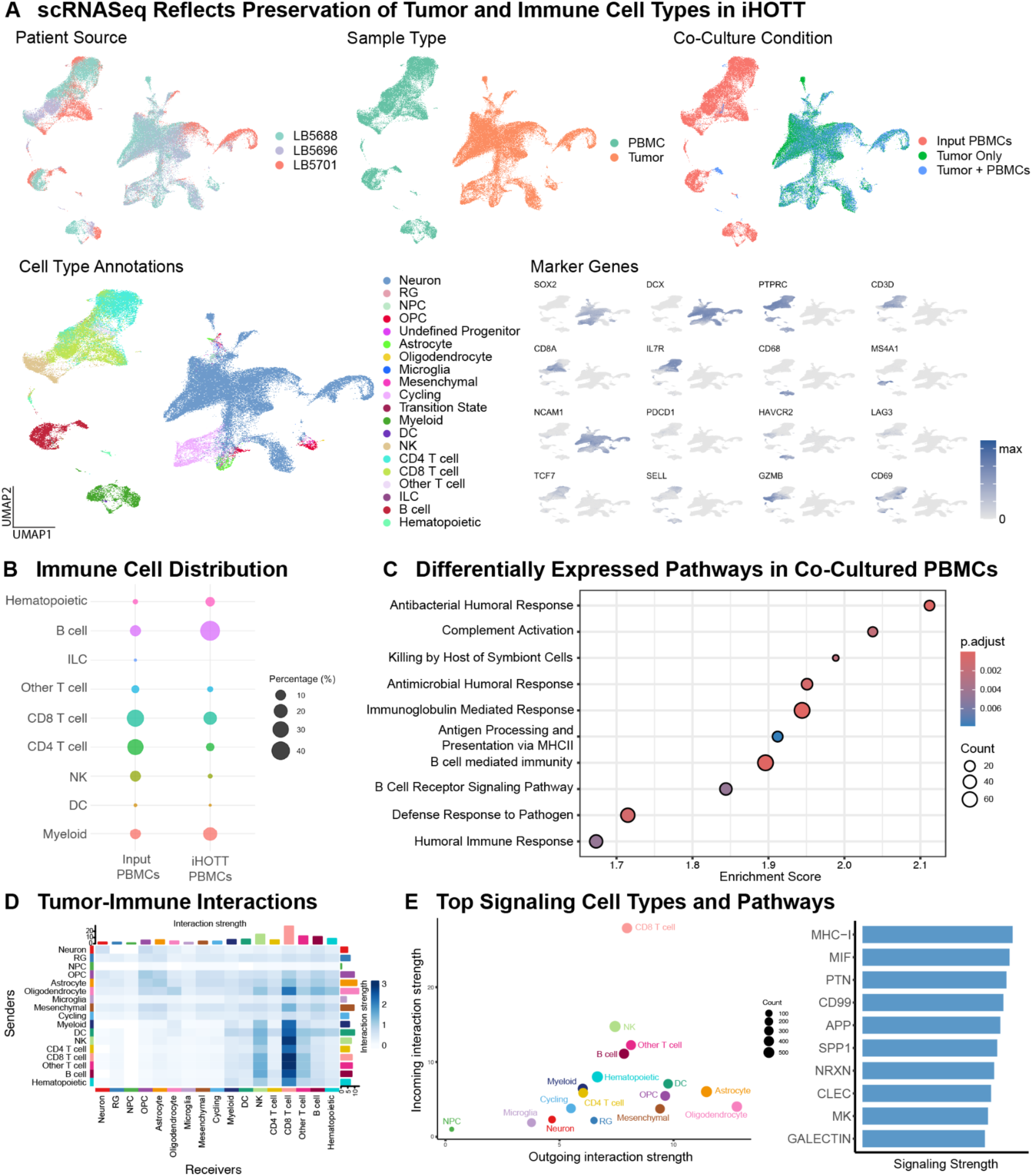
scRNA-seq Confirms Preservation of Tumor and Immune Compartments and Recapitulates Cell-Cell Interactions A) UMAPs from scRNA-seq of three patient samples comparing PBMC input, tumor-only cultures, and tumor + PBMC co-cultures. Top Left: cells annotated by patient source. Top Middle: cells annotated by tumor vs. PBMC origin. Top Right: cells annotated by co-culture condition. Bottom Left: Cells were annotated by reference projection onto the GBM meta-atlas and a PBMC reference object. Bottom Right: Feature plots show expression of representative marker genes validating tumor and immune cell type identities, activation, and exhaustion across conditions. B) Comparison of immune cell distributions between input PBMCs and PBMCs recovered from iHOTT after 7 days. All major immune populations, including CD4+ and CD8+ T cells, B cells, NK cells, and myeloid subsets, were preserved in co-culture. C) Gene Set Enrichment Analysis (GSEA) comparing input PBMCs and iHOTT PBMCs showed enrichment of immune activation pathways in the co-culture condition, suggesting engagement of functional immune programs. D) CellChat was used to analyze tumor and immune cell-cell interactions. We noted interactions between all cell types, with CD8 T cells as a notable receiver. E) CellChat analysis of ligand-receptor interactions between tumor and immune cells. Left: Heatmap of sender-receiver interactions showing CD8+ T cells as major communicators. Oligodendrocytes and astrocytes displayed strong outgoing signals. Right: Co-culture specifically induced upregulation of MHC-I and MIF signaling pathways, indicating active tumor-immune cross-talk within the iHOTT system.

Comparison of immune cell composition between input PBMCs and those recovered after co-culture in the iHOTT system demonstrated preservation of major immune cell types **[Figure 2B]**. Additionally, we sequenced four tumors that included a PBMC-only transplantation into the organoid in comparison with the matched PBMC input and the tumor-PBMC co-culture. Results showed that PBMC-only cultures were less effective at preserving NK cells and other T cell subsets, potentially reflecting reduced immune stimulation in the absence of tumor cells **[Supplementary Figure 4A-B].** While the composition observed in both datasets via single-cell sequencing differed slightly from our initial flow cytometry results, this variation likely reflects the increased specificity and resolution of using multiple markers for immune cell identification in the single-cell RNA sequencing analysis compared to single-marker analyses in flow cytometry.

A key feature of the iHOTT system is its ability to reveal how tumor populations are altered by exposure to immune cells. Compared to tumor-only conditions, co-culture with PBMCs resulted in shifts in tumor cell type composition, including expansions in progenitor populations and reductions in mesenchymal-like cells **[Supplementary Figure 5A]**. At the transcriptional level, tumor cells exhibited changes in gene expression, with enrichment of pathways related to metabolic function **[Supplementary Figure 5B]**.

Notably, the single-cell resolution of iHOTT enables precise interrogation of which cell types are responsible for secreting specific cytokines. Analysis of cytokine gene expression in the co-cultured condition revealed that certain growth factors, such as VEGFA, are predominantly expressed by tumor cells, whereas many chemokines—including CCL3, CCL4, and CXCL10— are primarily produced by myeloid and dendritic cell populations **[Supplementary Figure 5C]**.

### Interactions Between Tumor and Immune Cell Types are Preserved in iHOTT

Within the immune compartment of the three sequenced patients, we analyzed pathway enrichment in co-cultured immune cells relative to input PBMCs isolated directly from the patient. Several immune response pathways were significantly enriched in the iHOTT condition, including B cell–mediated immunity, complement activation, and antigen presentation **[Figure 2C]**. This upregulation reflects an activated immune response that emerges only in the presence of tumor cells within the culture system. Similar immune activation pathways have been observed in the glioblastoma tumor microenvironment and are associated with both endogenous and therapy-induced immune responses (28).

We further examined cell–cell interactions within our tumor–immune co-culture model. Notably, tumor oligodendrocyte-like cells emerged as significant signaling senders, while CD8⁺ T cells were the primary recipients of this signaling, followed closely by NK cells and other T cells (γδ T cells, MAIT cells, and double-negative T cells) **[Figure 2D-E]**. Among the most upregulated signaling pathways were MHC-I and macrophage migration inhibitory factor (MIF) **[Figure 2E].** MHC-I is essential for presenting intracellular antigens to CD8⁺ T cells, which aligns with the CD8 T cell’s central role in cytotoxic tumor surveillance, and MIF contributes to both immune cell recruitment and tumor immune evasion (29–32). The prominence of CD8⁺ T cells as signaling targets in this context is biologically consistent, as these cells depend on MHC-I-mediated antigen presentation to recognize and respond to tumor-derived antigens (33). Together, these results suggest that our autologous human co-culture model not only preserves the diversity of tumor and immune cell populations but also facilitates dynamic and physiologically relevant interactions that reflect the complex tumor–immune crosstalk occurring in patients.

### Pembrolizumab Treatment in iHOTT Results in Expansion of B and T Cells

Immune checkpoint inhibitors, particularly those targeting the PD-1/PD-L1 axis, have revolutionized cancer therapy by reactivating suppressed T cells and promoting immune cell expansion and function (34,35). Pembrolizumab is an anti-PD-1 monoclonal antibody that has been trialed as a checkpoint inhibitor in glioblastoma (36–38). Thus far, efficacy in glioblastoma has been limited, with trials reporting modest response rates in recurrent glioblastoma patients (6,36,37,39). We chose to evaluate the effect of pembrolizumab in our model, both to benchmark against our patient response data and to better understand the mechanism behind the limited response to pembrolizumab seen in patients.

Three patient-matched tumor and PBMC samples were co-cultured in our system in the presence of either an IgG control antibody or pembrolizumab over 7 days as previously described. Samples were sorted for scRNAseq. Media was additionally collected for cytokine profiling. Sequencing data demonstrated preservation of immune and tumor cell types as previously seen in our model across all conditions and patients **[Figure 3A, Supplementary Figure 6A]**.

**Figure 3:**
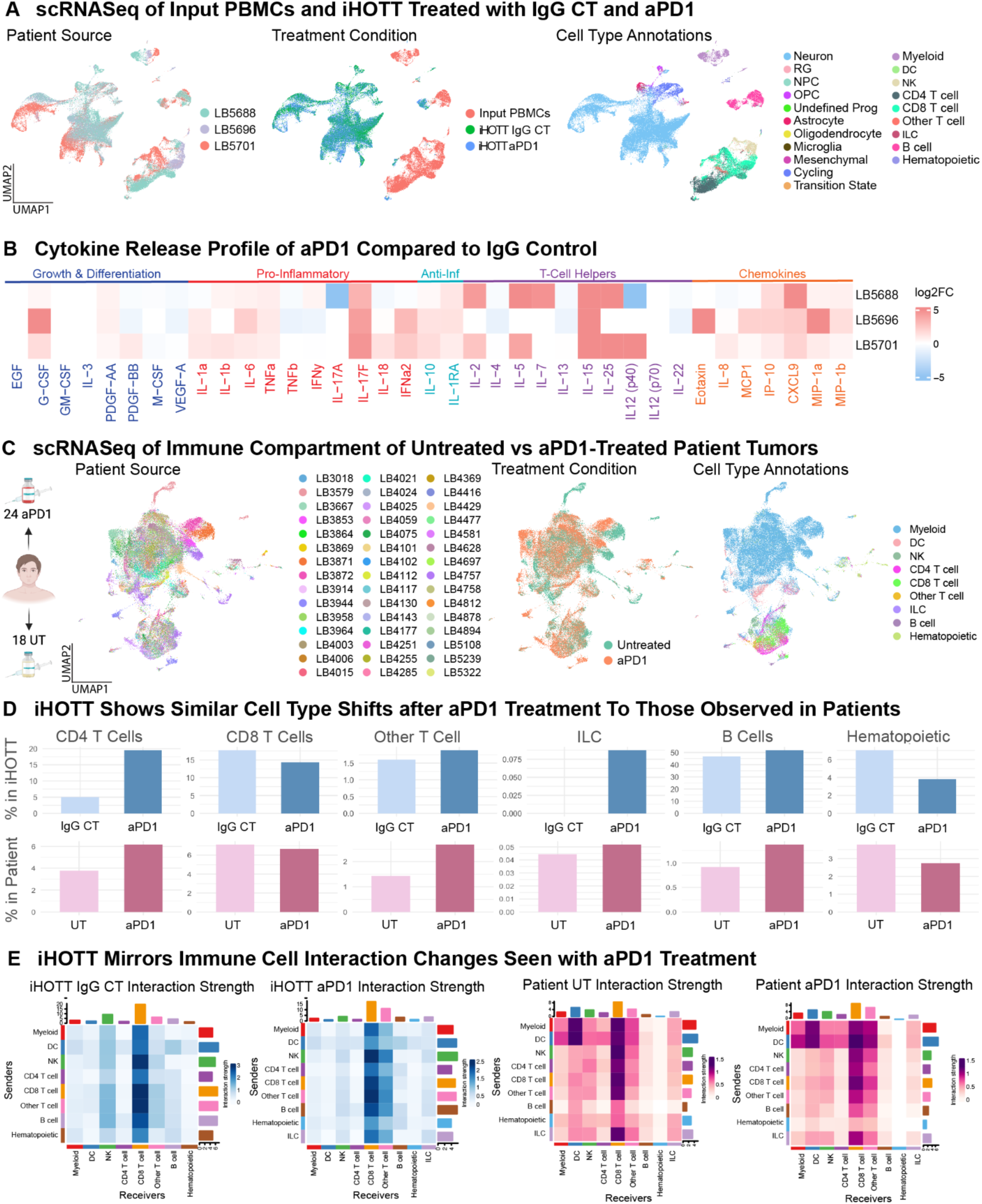
Pembrolizumab Treatment in iHOTT Recapitulates Cell Type and Interaction Changes Observed in Treated GBM Patients A) UMAPs from scRNA-seq of iHOTT samples annotated by patient source (left), treatment condition (middle), and cell type (right). All major tumor and immune cell populations were preserved across IgG control and pembrolizumab-treated samples. B) Cytokine profiling of iHOTT culture supernatants from three tumors treated with pembrolizumab or IgG control. Heatmap depicts relative cytokine expression in pembrolizumab-treated samples, with notable increases in IL-15, IL-25, and IL-17 following treatment. C) UMAPs from scRNA-seq of freshly isolated recurrent GBM patient tumor samples treated with either placebo or pembrolizumab. Left: annotated by patient source. Middle: annotated by treatment condition. Right: annotated by immune cell type, showing consistent detection of all major immune subsets observed in iHOTT, including CD4+ and CD8+ T cells, NK cells, B cells, and myeloid cells. D) Comparative analysis of cell type abundance post-treatment revealed parallel trends in iHOTT and patient samples, including increases in CD4+ T cells, B cells, innate lymphoid cells (ILCs), and non-canonical “Other T” populations in response to pembrolizumab. E) CellChat analysis of ligand-receptor communication networks post-treatment reflects that in iHOTT, pembrolizumab induced increased signaling activity involving ILCs and Other T cells (left). This was consistent with patterns observed in pembrolizumab-treated patient tumors (right). Strong interactions with CD8+ T cells and NK cells were maintained across all treatment conditions.

Further subclustering of immune cells in the scRNAseq dataset demonstrated that major cell types from the input PBMCs were retained in both treatment arms. Notably, modest increases in CD4⁺ T cells and B cells were observed in the pembrolizumab-treated condition **[Supplementary Figure 6B].** Analysis of differentially expressed pathways reflected a significant increase in immune response pathways in pembrolizumab-treated samples relative to input PBMCs, including pathways involved in B cell-mediated immunity, humoral immunity, and MHC II presentation **[Supplementary Figure 6C]**.

Subclustering of only the tumor cell fractions of the control and pembrolizumab-treated conditions reflected preservation of neuronal and mesenchymal cells with modest decreases in other tumor cell types after pembrolizumab treatment **[Supplementary Figure 6D].** Analysis of differentially expressed pathways showed an increase in biosynthesis pathways, synaptic pathways, and ribonucleotide processing **[Supplementary Figure 6E].** These findings suggest that checkpoint inhibition not only enhances immune activity but also reshapes the tumor cell transcriptional program, potentially remodeling the local microenvironment and altering tumor metabolic and signaling pathways in response to immune engagement.

### Pembrolizumab Treatment in iHOTT Alters Cytokine Signaling Profile

Cytokine profiling of iHOTT revealed key changes in response to pembrolizumab treatment **[Figure 3B, Supplementary Figure 7A-B].** Compared to IgG control, pembrolizumab-treated samples showed increased secretion of several cytokines, including IL-15, IL-17F, and IL-25 (IL-17E). IL-15 is a well-established driver of T and NK cell proliferation and survival, suggesting heightened cytotoxic lymphocyte activation in response to pembrolizumab treatment (40–42). IL-17F, typically produced by Th17 cells, is associated with inflammatory responses and may reflect broader immune engagement in the tumor microenvironment (43). IL-25, also known as IL-17E, plays a role in shaping type 2 immune responses and may contribute to lymphocyte recruitment or polarization (44). Together, these shifts in secreted cytokines indicate heightened immune engagement and lymphocyte activation in response to PD-1 blockade.

### iHOTT Mirrors Immune Cell Type Shifts Observed in Treated Patient Samples

Forty-two patients with recurrent glioblastoma at our institution had been previously enrolled in a randomized trial comparing pembrolizumab treatment versus no treatment on their first or second recurrence [*Supplementary Table 2*]. The immune fraction of these patient tumors was isolated and submitted for scRNAseq. Initial analysis confirmed the preservation of all major immune cell types, with a predominance of myeloid cells **[Figure 3C]**. This myeloid cell enrichment is consistent with previously published data regarding the composition of tumor-infiltrating lymphocytes and likely reflects the fact that these samples were sequenced immediately upon isolation, without any in vitro culture or expansion (45).

Notably, analysis of immune cell type distributions in pembrolizumab-treated versus untreated patients closely mirrored the shifts observed in the iHOTT model following pembrolizumab exposure **[Figure 3D]**. This concordance supports the physiological relevance of iHOTT as a platform for modeling autologous immune responses to checkpoint blockade and suggests that the observed treatment-induced changes are reflective of patient biology.

### iHOTT Mirrors Shifts in Cell-Cell Interactions Observed in Treated Patient Samples

Cell-cell interaction analyses were performed in iHOTT IgG control and pembrolizumab-treated samples, as well as in patient untreated and pembrolizumab-treated samples. As previously described, co-cultured samples in iHOTT demonstrated multiple interactions between tumor and immune samples **[Supplementary Figure 8A].** For further analysis, samples were subclustered to immune cell types only. All samples showed significant interactions with CD8 T cells. Furthermore, the increase in interactions going to other T cells (γδ T cells, MAIT cells, and double-negative T cells) seen in treated patients was mirrored in iHOTT **[Figure 3E]**. While the role of CD8⁺ T cells as a key receiver of immune signaling is established, recent studies suggest the possibility that unconventional T cell populations, such as γδ T cells and MAIT cells, may also be activated or expanded in response to checkpoint inhibition (46).

Overall pathways enriched in iHOTT pembrolizumab-treated samples included resistin, WNT, complement, and CSF, all key immune activation pathways **[Supplementary Figure 8B]**. When samples were subclustered to only examine immune cells, we observed that signaling pathways involved in immune responses reflected similar trends between untreated and treated samples in both patients and iHOTT **[Supplementary Figure 9A-C].** For example, CD45 signaling was relatively enriched in untreated samples, whereas complement, CD226, TIGIT, and CD70 were upregulated in pembrolizumab-treated samples, suggesting enhanced co-stimulatory signaling, immune cell activation, and potential cytotoxic engagement in response to checkpoint blockade (47). CD226 and TIGIT are part of the TIGIT-CD226-CD96 checkpoint axis, and their upregulation may reflect activation of this axis, which is known to regulate the balance between T cell activation and inhibition through shared ligand competition (48). Together, these findings reinforce the ability of iHOTT to capture complex, clinically relevant immunologic responses to PD-1 inhibition.

In addition, cell types involved in selected pathways were consistent between iHOTT and patient samples **[Supplementary Figure 9C].** These parallel shifts in cell type involvement reinforce the relevance of iHOTT in capturing key aspects of the immune response to PD-1 inhibition observed in patients.

### T Cell Receptor Sequencing (TCRseq) in iHOTT Reveals an Increase in Clonotype Diversity Due to the Production of Novel Clones

T cell receptors (TCRs) are highly diverse surface proteins expressed on T cells that enable antigen recognition and immune specificity. Each T cell expresses a unique TCR formed by somatic recombination of variable (V), diversity (D), and joining (J) gene segments, producing an immense repertoire of T cells capable of recognizing a wide array of antigens (49). TCR sequencing allows for high-resolution profiling of T cell clonotypes (shared TCR sequences in a population), offering insight into the clonal dynamics of an immune response. In the context of antigen stimulation, an effective T cell response is typically characterized by the expansion of a limited number of dominant clonotypes targeting specific antigens (50–52).

Given that pembrolizumab treatment in our model led to a clear expansion of T cells and that many of the observed tumor–immune interactions involved T cell populations, we sought to further investigate the nature of the T cell response by performing TCR sequencing across both patient and iHOTT samples.

TCR sequencing of input PBMCs, IgG control co-cultures, and pembrolizumab-treated co-cultures revealed progressive shifts in clonal composition in response to tumor exposure and immune checkpoint blockade. The proportion of unique clonotypes increased stepwise following tumor co-culture and further with pembrolizumab treatment, indicating that both antigenic stimulation and therapeutic modulation contribute to diversification of the T cell repertoire. Treated samples also exhibited a greater proportion of rare clonotypes, suggesting expansion or recruitment of previously low-frequency populations **[Figure 4A]**. These trends were further reflected by alterations in TCR chain length distributions and an increase in overall diversity index in pembrolizumab-treated samples, consistent with more dynamic and heterogeneous clonal architecture **[Supplementary Figure 10A-B]**.

**Figure 4:**
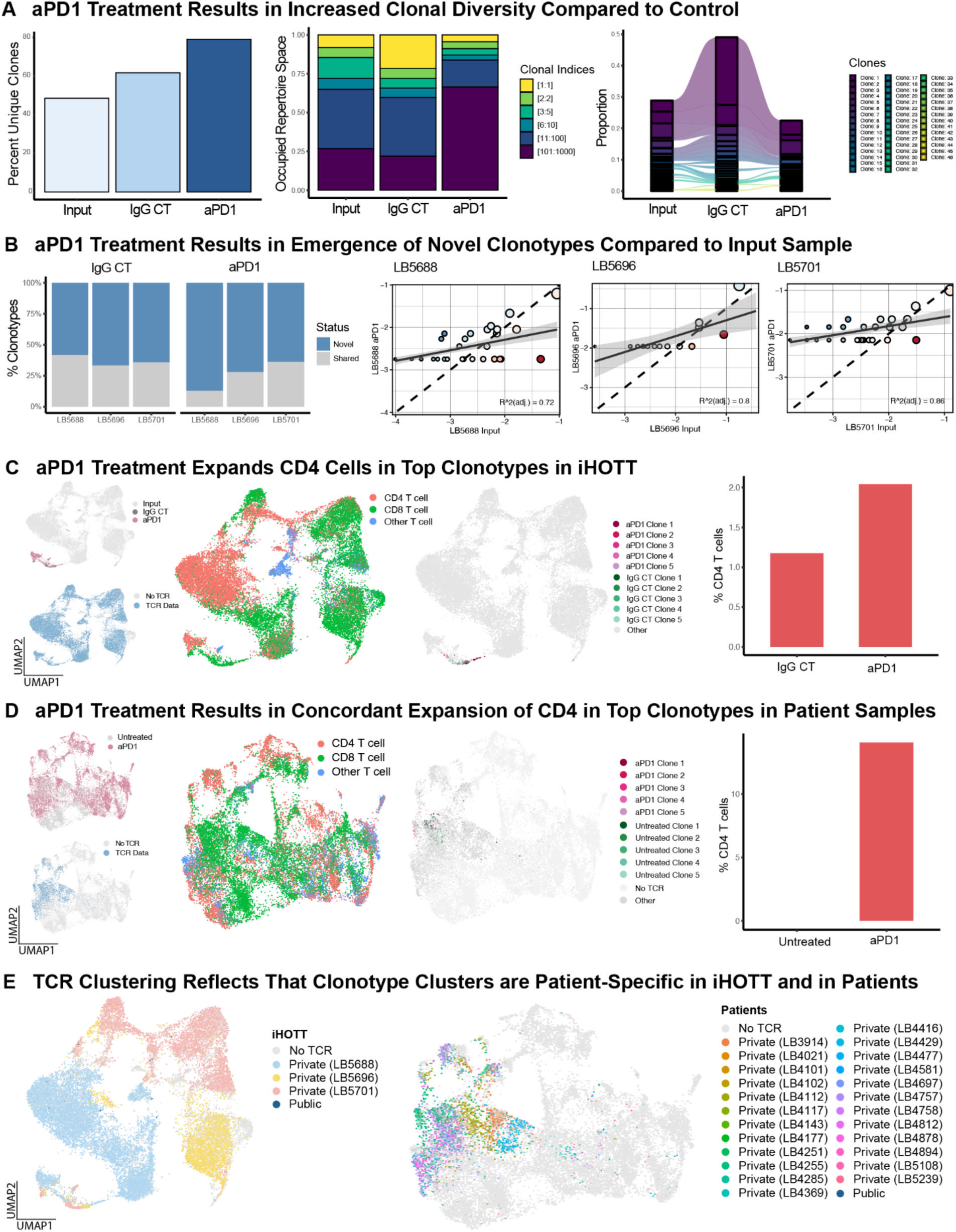
TCR Sequencing Reveals Pembrolizumab-Induced Clonal Remodeling of T Cells in iHOTT and Patient Samples A) TCR sequencing of iHOTT samples across three patients. Left: Quantification of unique TCR clonotypes revealed a stepwise increase in clonal diversity from input PBMCs to IgG control, and a further increase in pembrolizumab-treated samples. Middle: Repertoire occupancy analysis showed that pembrolizumab treatment resulted in greater representation of rare clonotypes, shifting the overall clonal architecture. Right: Comparison of the top 20 clones in each condition demonstrated the emergence of new clonotypes in pembrolizumab and control samples, alongside persistence of some input-derived dominant clones. B) Left: Assessment of novel clonotype composition showed a significant increase in the proportion of the repertoire composed of new clonotypes in pembrolizumab-treated samples compared to both input and control conditions. Right: Scatter plot comparing clone frequencies between pembrolizumab-treated and input samples showed emergence of new clones and partial overlap with the input repertoire. C) Left: UMAPs of reclustered T cells from iHOTT showing treatment condition (input, IgG control, pembrolizumab) and TCR detection. All samples retained robust TCR coverage. Middle: Cells labeled by treatment and by the top 5 expanded clonotypes within control and pembrolizumab conditions. Right: Top clonotypes in the pembrolizumab condition were enriched for CD4+ T cells, suggesting clonal expansion and reinvigoration mediated by CD4+ populations following PD-1 blockade. D) Analogous TCR analysis in recurrent GBM patient samples treated with pembrolizumab or placebo. While TCR coverage was approximately 30%, consistent trends were observed. Pembrolizumab-treated tumors demonstrated the emergence of novel clonotypes, with top clones reflecting an increase in CD4+ T cells. E) GLIPH2 analysis of TCR sequences was used to group clonotypes based on predicted shared antigen specificity. In both iHOTT (left) and patient samples (right), clonotype clusters were predominantly patient-specific, with limited sharing across individuals. This suggests antigenic heterogeneity in GBM.

To determine the contribution of novel versus persistent clonotypes to clonal diversity, we compared TCR repertoires across treatment conditions. We refer to’novel’ clonotypes here as those not detected in the input PBMCs but were observed following co-culture or treatment, reflecting likely expansion from rare precursors. Approximately half of the clonotypes present in the IgG control conditions were novel across tumors compared to the matched input, representing the expansion of novel T cell clones in response to the tumor. This effect was amplified in the pembrolizumab-treated group, which exhibited an increased number of novel clonotypes, likely reflecting heightened immune activation or an enhanced immune response driven by checkpoint inhibition. **[Figure 4B]**. When analyzed individually, each pembrolizumab-treated sample showed evidence of novel clonotype expansion compared to the input PBMCs. However, the overlap between clonotypes in the pembrolizumab-treated samples and the input PBMCs was comparable to that observed in the IgG control condition, suggesting that the observed increase in clonal diversity is primarily driven by the expansion of newly detected clonotypes **[Supplementary Figure 10C].**

### Integration of TCR Clonotypes with Transcriptional Profiles Reveals CD4-Driven Response to Pembrolizumab Treatment in iHOTT and in Patients

To further investigate the cellular and transcriptional features of dominant T cell clonotypes, we integrated TCR sequencing data with our single-cell transcriptomic dataset. The PBMC-only subset was filtered to include T cells and re-clustered to specifically examine T cell-specific changes. The dataset was annotated as previously described, and marker genes were visualized for T cell subtypes, T cell progenitors, activation, and exhaustion **[Supplementary Figure 10D]**. TCR coverage analysis showed that approximately 45% of our T cells had clonotype information. We then identified the five most expanded clonotypes, each composed of approximately 4 to 54 cells, and visualized their distribution across the T cell UMAP. Clonal composition analysis revealed that dominant clones were associated with CD4 T cell expansion **[Figure 4C]**.

A separate analysis was performed to distinguish novel from persistent clonotypes across treatment conditions. In pembrolizumab-treated co-cultures, novel clonotypes were significantly enriched for CD4 T cells compared to control samples **[Supplementary Figure 10E]**. Further analysis was performed for marker expression within T cells stratified by treatment condition. Results showed an increase in TCF7 and LEF1 expression as well as a reduction in TOX expression after checkpoint inhibition **[Supplementary Figure 10F]**. This gene expression pattern is consistent with a stem-like precursor phenotype that has been associated with effective responses to checkpoint blockade (18,55). Prior studies have shown that *TCF7*⁺ CD4 and CD8 T cells function as progenitor populations capable of self-renewal and differentiation into functional effector cells following PD-1 inhibition (18,55). These findings suggest that pembrolizumab promotes the maintenance of stem-like CD4 T cell populations within the co-culture system.

### Checkpoint Inhibition Responses in Patients Mirror iHOTT and Reflect Personalized Clonal Landscapes

TCR sequencing data from untreated and pembrolizumab-treated patient samples were aligned with matched single-cell transcriptomic profiles, and a focused analysis was performed on the T cell compartment. The proportion of unique clonotypes and the overall clonotype distribution were assessed across samples. This analysis revealed a heterogeneous pattern among patients, with no consistent trends distinguishing untreated from treated conditions, highlighting the variability of clonal responses to pembrolizumab at the patient level **[Supplementary Figure 11A-B].**

To further assess patient-level changes following treatment, the dataset was filtered and re-clustered to include only T cells. Approximately 16% of cells were found to have paired TCR information. The top five clonotypes within each treatment condition were identified, with each clonotype comprising 10 to 33 cells. In line with observations from the iHOTT model, CD4 cells were expanded in the top clonotypes **[Figure 4D, Supplementary Figure 11C]**. This concordance between patient samples and the iHOTT model underscores the physiological relevance of the system and its ability to recapitulate key aspects of human immune responses to immunotherapy.

Finally, the potential for shared antigen specificity among clonotypes across individuals was assessed using GLIPH2 (Grouping of Lymphocyte Interactions by Paratope Hotspots 2), a computational algorithm that clusters T cell receptors based on predicted common antigen recognition despite sequence variability (56). Clusters were classified as “private” when composed of T cells from a single individual, or “public” when containing T cells from multiple individuals, indicative of convergent antigen targeting. This analysis was conducted across both the iHOTT model and patient datasets. While a limited number of public clonotypes were identified in iHOTT, the majority of clusters across both iHOTT and patients were private and restricted to individual patients **[Figure 4E]**. These results suggest that the T cell response to pembrolizumab in glioblastoma is largely patient-specific, with minimal evidence of convergence onto shared tumor antigens. This highly individualized response likely contributes to the limited clinical efficacy of PD-1 blockade in GBM, as heterogeneous immune recognition across patients would inherently prevent convergence toward a broadly effective therapeutic response, underscoring the necessity for more targeted immunotherapy strategies. Notably, the iHOTT model recapitulated this largely private, individualized clonal architecture, further supporting its utility as a physiologically relevant platform for modeling patient-specific immune responses to immunotherapy.

## DISCUSSION

In this study, we present iHOTT as a human-specific, autologous organoid model that enables detailed evaluation of tumor–immune interactions in glioblastoma. Through multimodal characterization, including cell type composition analyses, flow cytometry, cytokine profiling, cell-cell interaction inference, and TCR clonotyping, we demonstrate that iHOTT closely benchmarks to patient tumor samples and recapitulates patient-specific responses to pembrolizumab treatment. The preservation of key immune and tumor compartments, along with the emergence of treatment-induced features that parallel clinical samples, supports the physiological relevance of this system.

Patient-matched PBMCs were intentionally selected to model the early immune response to glioblastoma, simulating the initial infiltration of circulating immune cells into the tumor microenvironment. Unlike tumor-infiltrating lymphocytes, which may exhibit features of chronic activation or exhaustion, PBMCs represent a responsive immune population encountering tumor antigens. This design allows for the interrogation of early immune activation, tumor recognition, and response to immunotherapeutic modulation in a context that reflects the dynamics of immune entry in patients. In the future, it may be possible to extend this system to model the chronic inflammatory conditions observed in tumor-infiltrating lymphocytes. For example, options include prolonging culture duration, incorporating myeloid-driven immunosuppressive signals, or reconditioning PBMCs ex vivo to enable investigation of both early and late-stage immune responses within the same platform.

Although major lymphocyte populations were preserved, we noted a relative underrepresentation of myeloid cells in the iHOTT system compared to fresh patient tumor samples. This discrepancy likely reflects limited myeloid cell survival or retention in culture, a challenge commonly observed in in vitro models due to the short lifespan and high plasticity of monocytes and dendritic cells ex vivo. The absence of tissue-derived stromal or vascular cues may also limit myeloid cell recruitment and maintenance. This highlights an important area for future optimization, particularly given the central role of myeloid populations in glioblastoma immunosuppression, antigen presentation, and T cell regulation.

Potential strategies to enhance myeloid representation include media supplementation with cytokines such as GM-CSF, IL-4, or M-CSF to support differentiation and survival or additional chemokine modulation. Extending co-culture duration or including patient-derived microglia and macrophages may also help capture the full spectrum of tumor-associated myeloid diversity. Addressing this limitation will be crucial for leveraging iHOTT to study immunosuppressive circuits and evaluate therapeutic interventions targeting the myeloid compartment in glioblastoma.

Despite this limitation, iHOTT is particularly well-suited for studying T cell–mediated and CAR-T– based immunotherapies. The system provides a human-specific, autologous platform that preserves patient-matched tumor and immune compartments, allowing for highly relevant modeling of tumor-T cell interactions. By enabling precise tracking of clonal dynamics through TCR sequencing, iHOTT allows researchers to identify expanding or contracting clonotypes in response to tumor antigens or immunotherapy. It also permits assessment of antigen specificity and T cell functional states, including activation, exhaustion, and stem-like phenotypes, at single-cell resolution.

This is especially valuable for evaluating CAR-T efficacy in glioblastoma, a setting where immunosuppressive barriers, antigen heterogeneity, and limited T cell infiltration have hampered clinical translation. In iHOTT, CAR-T cells can be introduced into a native tumor microenvironment and monitored for proliferation, cytotoxicity, and engagement with target antigens. Because the system supports multimodal analysis, including flow cytometry, cytokine profiling, and transcriptional mapping, it enables comprehensive mechanistic dissection of T cell behavior in a way that static endpoint assays cannot achieve.

Beyond mechanistic discovery, iHOTT holds strong promise for translational applications including preclinical drug screening, immune engineering, and precision immunotherapy development. Its capacity to recapitulate inter-patient variability, including individualized clonotype usage and immune responses, supports its use as a personalized testing platform to predict therapeutic efficacy or resistance. By capturing key cellular, transcriptional, and clonal features of the human anti-tumor immune response, iHOTT offers a powerful and versatile tool to bridge the gap between bench research and individualized immunotherapeutic strategies in glioblastoma.

## AUTHOR CONTRIBUTIONS

Conceptualization - AB, SB, EF

Methodology - SB, EF, AB

Validation - SB, EF, AB, ML, AM, DA

Formal Analysis - SB, EF, AB, ML, AM, DA

Investigation - SB, EF, AB, ML, AM, DA, LS, IP, AM, GH

Resources - AB, RP, DN, LL, KP, RE, WK, AW

Data Curation - SB, EF, AB

Writing - Original Draft - SB, AB

Writing - Revision & Editing - All authors

Visualization - SB, EF, AB

Supervision - AB

Project Administration - AB

Funding Acquisition - AB (*Additional fellowships in acknowledgments)*

## ACKNOWLEDGMENTS

We would like to thank the members of the Bhaduri Lab for their insightful advice and comments on the study. We thank the Broad Stem Cell Research Center Flow Cytometry Core, the Virology Core, and the Immune Assessment Core at UCLA. We would further like to thank Suhua Feng for his help with the sequencing of our samples. We would like to thank Su Aung, Sergey Mareninov, and Adrian Murrillo at the Brain Tumor Translational Research Core (BTTR) for enabling tumor sample acquisition. We would like to thank Marc Perry and Maximilian Haeussler (UCSC) for their help in compiling the CellBrowser.

This study was generously funded by support to AB from: Swim Across America, Jonsson Comprehensive Cancer Center, Sloan Research Fellowship from the Alfred P. Sloan Foundation, NIH NCI P50CA211015 including a Career Enhancement Program Award, The Sontag Foundation (Distinguished Scholar Award), V Scholar Award from The V Foundation, The Uncle Kory Foundation, The American Cancer Society (CSCC-Team-23-980262-01-CSCC), The Margaret Early Medical Research Trust, The Pew Foundation and The Alexander and Margaret Stewart Trust, McKnight Neurobiology of Brain Disorders Award (with KP), The Rose Hills Foundation and the Broad Stem Cell Research Center.

Additional funding was provided to SB by the Tumor Cell Biology T32 at UCLA and the Blalock Foundation H&H Lee Resident Research Grant. Funding for EF was provided by the David Geffen Scholarship and the UCLA-Caltech Medical Scientist Training Program (T32GM152342).

## DATA AVAILABILITY

The original datasets generated here are viewable on an interactive public UCSC Cell Browser. Data is available under these links:

https://cells-test.gi.ucsc.edu/?ds=ihott https://cells-test.gi.ucsc.edu/?ds=ihott+untreated https://cells-test.gi.ucsc.edu/?ds=ihott+treated

The data are also available as Seurat objects for download. The raw data collected from this study are being deposited in dbGAP. The reference patient dataset used for benchmarking our results was previously published (57,58).

## METHODS

### Cell & Tissue Culture

This study involved the use of multiple human-derived components, including primary tumor tissue, peripheral blood mononuclear cells (PBMCs), cortical organoids, and gliomasphere cultures. Detailed protocols for each of these components are described in the relevant sections below. Briefly, primary glioblastoma tumor tissue and matched peripheral blood were obtained intraoperatively from consented patients under IRB-approved protocols (see *Patient Tissue Acquisition*). Cortical organoids were generated from human embryonic stem cells (UCLA6 line) and matured for use in co-culture experiments as detailed in *Cortical Organoid Generation*.

### Gliomasphere Culture

This study also utilized patient-derived gliomasphere lines (generously provided by the Nathanson Lab at UCLA) for the optimization experiments before the utilization of fresh patient tissue. Gliomaspheres were cultured in Neurosphere Media composed of DMEM/F12 (ThermoFisher cat# 11330057), 5 mL of 100x Glutamax (ThermoFisher cat# 35050061), 5 mL of 100x Pen-Strep (ThermoFisher cat# 15140122), and 10 mL of 50x B27 Supplement without Vitamin A (ThermoFisher cat# 12587010) per 500 mL bottle. Cultures were supplemented every 5 days with 1x HEF (2.5 µL of 400x stock per mL of media), a mixture containing 5 µg/mL heparin, 20 ng/mL FGFb (ThermoFisher cat# PHG0263), and 50 ng/mL EGF (ThermoFisher cat# PHG0313).

Cells were maintained in non-adherent, filter-cap cell culture–treated flasks at 37°C with 5% CO₂. Culture density was kept between 50,000–200,000 cells/mL, with an optimal range of 100,000 cells/mL. Spheres were passaged every 4–10 days based on growth rate, sphere size, and media color; a 7-day interval was typically sufficient. Cells were dissociated using TrypLE Express (ThermoFisher cat# 12605028) and frozen in Bambanker freezing media (Bulldog Bio cat# BB01) when needed.

### Patient Tissue Acquisition

Primary glioblastoma specimens and matched peripheral blood were collected from patients undergoing surgery at Ronald Reagan UCLA Medical Center under IRB-approved protocols (IRB #10-000655 and #21-000108), following informed research consent. Samples were processed immediately following surgical resection for either tumor dissociation or PBMC isolation as described below.

### Tumor Cell Dissociation

Tumor samples were obtained fresh from the operating room and were processed on the same day. Tissue was first manually dissected into small fragments using a sterile scalpel. The fragments were then transferred to 5 mL tubes containing 2.5 mL of Papain and 125 µL of DNase I prepared per manufacturer’s protocols (Worthington). Samples were incubated at 37°C for 45– 60 minutes, with intermittent manual agitation every 5-10 minutes to promote tissue breakdown.

Following incubation, tissues were gently triturated and centrifuged at 300 × g for 5 minutes to isolate a pellet of single cells. The papain-containing supernatant was removed, and the cell pellet was resuspended in Sasai3 media (recipe described above in *Organoid Preparation*).

Tumor cell suspensions were passed through a 40 µm cell strainer to remove debris and counted using a hemocytometer. For GFP labeling, 5 million tumor cells were resuspended in 1.5 mL total volume in Eppendorf tubes containing 1 mL Sasai3 media, 1 µL polybrene (final concentration 8 µg/mL), and 2 µL of CMV-GFP lentivirus (SignaGen) per 1 million cells. Tubes were incubated on a rotator at 37°C for 90 minutes to allow transduction. After incubation, cells were washed three times with PBS to remove excess virus and polybrene, and the final pellet was resuspended in fresh media for immediate use in organoid co-culture experiments, as described in the *iHOTT Model Generation and Culture* section.

### PBMC Isolation

Peripheral blood was collected intraoperatively from patients and processed fresh on the day of surgery. Blood was diluted up to 50 mL with sterile PBS and gently mixed by pipetting. The diluted blood was then carefully layered over 20 mL of Ficoll-Paque PLUS (GE/Cytiva) in two separate 50 mL conical tubes (10 mL per tube), allowing the blood to drip slowly down the side to avoid disturbing the interface. Samples were centrifuged at 400 × g for 30 minutes at 18°C with no brake on the centrifuge.

The PBMC layer typically appears as a white, cloudy interface after centrifugation. It was extracted using a sterile plastic Pasteur pipette by gently aspirating just above the Ficoll layer. Extracted cells were transferred to a clean 50 mL tube and topped off with PBS. Samples were centrifuged at 400 × g for 10 minutes at 4°C. The supernatant was poured off, and cells were resuspended in 10 mL of PBS and centrifuged again under the same conditions. Cells were then counted using Trypan Blue exclusion and a hemocytometer.

PBMCs were either used fresh for transplant, and any remaining PBMCs were cryopreserved. For cryopreservation, cells were resuspended at 5–20 million cells/mL in freezing media composed of 90% heat-inactivated fetal bovine serum (FBS; filtered) and 10% DMSO. 1 mL of cell suspension was aliquoted per cryovial and frozen at –80°C.

### Cortical Organoid Generation

Human embryonic stem cells (hESCs, UCLA6 line) were maintained under feeder-free conditions in Matrigel-coated 6-well plates using mTeSR Plus medium (STEMCELL Technologies) supplemented with 10% mTeSR Plus Supplement and 1× Primocin (InvivoGen). Media was replaced every other day, and cells were passaged when cultures reached approximately 75– 90% confluence.

For passaging, ReLeSR (STEMCELL Technologies) was added to each well and incubated at room temperature for 1 minute before aspiration. Plates were then incubated at 37°C for an additional 5 minutes to facilitate detachment. Cells were gently disaggregated into small clusters and replated at a 1:4 to 1:6 ratio onto fresh Matrigel-coated plates. For cryopreservation, passaged cells were resuspended in mFreSR freezing media (STEMCELL Technologies), aliquoted into cryovials, and frozen at –80°C for 24–48 hours before transfer into liquid nitrogen for long-term storage.

Cortical organoids were generated as previously described (27). Briefly, hESCs at ∼80% confluence were incubated with 1 mL Accutase (STEMCELL Technologies) per well for 5 minutes at 37°C. Detached cells were collected, centrifuged at 300 × g for 5 minutes, and resuspended in Sasai1 medium, composed of GMEM (Gibco), 20% KnockOut Serum Replacement (ThermoFisher), 0.1 mM β-mercaptoethanol, 1× Non-Essential Amino Acids (NEAA), 1× Sodium Pyruvate, and 1× Primocin.

Cells were seeded at a density of 1 million cells divided evenly across a 96-well low-attachment V-bottom plate in Sasai1 medium. Sasai1 medium consisted of GMEM (ThermoFisher, cat# 11710-035), 20% KnockOut Serum Replacement (ThermoFisher, cat# 10828-028), 0.1 mM β-mercaptoethanol (Sigma, cat# M3148), 1× Non-Essential Amino Acids (CCF), 1× Sodium Pyruvate (CCF, 11 mg/mL stock), and 1× Primocin (CCF). This media was further supplemented with 20 μM Y-27632 (ROCK inhibitor), 5 μM SB431542 (TGF-β receptor inhibitor), and 3 μM IWR-1-endo (Wnt pathway inhibitor) to promote aggregation and neuroectodermal induction. Aggregates were cultured for 72 hours undisturbed, after which half the media was replaced with fresh Sasai1 containing the same inhibitors. Media was then changed every other day until Day 7, with continued supplementation with inhibitors, after which Y-27632 was omitted from the medium.

On Day 18, organoids were transferred to ultra-low attachment 6-well plates and cultured in Sasai2 medium, composed of DMEM/F12 with Glutamax (ThermoFisher, cat# 10565-018), 1× N-2 Supplement (ThermoFisher, cat# 17502-048), Lipid Concentrate (ThermoFisher, cat# 11905-031), and 1× Primocin. From Day 35 onward, organoids were cultured in Sasai3 medium consisting of DMEM/F12 with Glutamax (cat# 10565-018), 1× N-2 Supplement (cat# 17502-048), Lipid Concentrate (cat# 11905-031), 10% FBS (HyClone, cat# SH30071.03), 5 μg/mL heparin (Sigma, cat# H3149), 0.5% vol/vol growth factor–reduced Matrigel (BD Biosciences, cat# 354230), and 1x Primocin. Media changes were performed every other day throughout the culture period.

### iHOTT Model Generation and Culture

iHOTT co-cultures were established using mature cortical organoids (weeks 8–12) generated as described above in *Cortical Organoid Generation*. Freshly dissociated primary tumor cells and matched PBMCs were prepared on the same day, following protocols outlined in the *Tumor Cell Dissociation* and *PBMC Isolation* sections.

For co-culture, 500,000 GFP-tagged tumor cells and 500,000 PBMCs (1:1 ratio) were combined and transplanted onto organoids using a hanging drop method. Organoids were transferred to the inverted lid of a 10 cm dish using wide-bore pipette tips, and excess media was removed. A 10 μL droplet containing tumor cells and PBMCs was gently placed atop each organoid. The lid was then inverted over a 10 cm dish containing 10 mL of base culture media to maintain humidity and prevent evaporation. Hanging drop co-cultures were incubated at 37°C for 12–16 hours before transfer to ultra-low attachment 6-well plates in Sasai3 medium. Tumor cells and PBMCs typically surrounded the organoid and began migrating inward over the following days.

Organoid co-culture models were then cultured for 7 days in 50-50 Sasai3:RPMI media without additional supplementation. Cultures were maintained under continuous rotation on an orbital shaker with media changes every other day. On day 7, organoids were harvested for downstream analysis, including immunofluorescence, flow cytometry, single-cell RNA sequencing, cytokine profiling, and TCR sequencing, as detailed in subsequent sections.

### Immunofluorescence

Organoids were fixed in 4% paraformaldehyde (PFA) at room temperature for 45 minutes, followed by three washes with PBS. Fixed organoids were incubated overnight in 30% sucrose at 4°C, embedded in OCT compound over dry ice, and stored at –80°C. Cryosections (12 µm thick) were collected on Superfrost Plus slides and air-dried for at least 1 hour at room temperature prior to storage and staining.

Sections were washed in PBS. Antigen retrieval was performed with a boiled citrate-based antigen retrieval solution for 20 minutes. Samples were then permeabilized and blocked with blocking buffer containing 5% donkey serum, 3% bovine serum albumin (BSA), and 0.1% Triton-X in PBS for 1 hour at room temperature. Primary antibodies were diluted 1:200 in blocking buffer and incubated overnight at 4°C in a humidified chamber. The following primary antibodies were used:

● **CD3 epsilon** (rabbit, Abcam, cat# ab16669)
● **CD14** (mouse, Abcam, cat# ab181470)
● **CD19** (rabbit, Cell Signaling Technology, cat# 3574S)
● **GFP** (goat, Novus Bio, cat# NB100-1770)

Following primary incubation, sections were washed three times with PBS containing 0.1% Triton-X and incubated with species-specific secondary antibodies for 2 hours at room temperature in the dark. The secondary antibody mix also included DAPI (Thermo Fisher) at a 1:1000 dilution. All secondary antibodies were diluted 1:1000 in blocking buffer and included:

● **Alexa Fluor 488 anti-goat (donkey host)**, Invitrogen, cat# A32814
● **Alexa Fluor 647 anti-mouse (donkey host)**, Invitrogen, cat# A32787
● **Alexa Fluor 647 anti-rabbit (donkey host)**, Invitrogen, cat# A31573

Nuclei were counterstained using DAPI (ThermoFisher) for 5 minutes before a final PBS wash. Slides were mounted with ProLong Gold Antifade Mountant (ThermoFisher) and stored at 4°C prior to imaging.

Imaging was performed on an EVOS cell imaging system. Identical exposure and acquisition settings were applied across samples for any experiments involving quantification. Representative fields were selected for display.

### Dissociation, Sorting, and Flow Cytometry

Organoids were harvested on day 7 of co-culture and dissociated into single-cell suspensions using enzymatic digestion. For each sample, tumor-transplanted organoids were transferred to a 1.7 mL Eppendorf tube containing 1 mL of Papain solution and 50 μL of DNase I (Worthington), then incubated at 37°C for approximately 30 minutes. To enhance tissue breakdown, samples were shaken vigorously by hand for 10 seconds every 5 minutes during incubation. After enzymatic digestion, samples were triturated with a P1000 pipette and centrifuged at 300 × g for 5 minutes. The resulting cell pellet was resuspended in cold FACS buffer (PBS with 2% FBS and 0.5 mM EDTA), filtered through a 40 µm strainer, and stained with DAPI (1 µg/mL) for viability assessment.

For **FACS sorting**, single-cell suspensions were analyzed on a BD FACSAria machine. Cells were gated to exclude debris, doublets, and dead cells using forward/side scatter and DAPI exclusion. GFP+ tumor cells and CD45+ immune cells were sorted for downstream applications, including single-cell RNA sequencing. GFP gating thresholds were set using non-transplanted control organoids as negative controls.

In parallel, **flow cytometric profiling** of dissociated iHOTT samples was performed using an Attune NxT Flow Cytometer (ThermoFisher). Cells were stained with a panel of fluorophore-conjugated antibodies to define immune subpopulations. The following antibodies were used:

● **CD3** (AF700, clone UCHT1, ThermoFisher, cat# 56-0038-80)
● **CD14** (APC, clone M5E2, BioLegend, cat# 301807)
● **CD19** (APC-eFluor780, clone HIB19, ThermoFisher, cat# 47-0199-41)
● **CD45** (PE, clone HI30, ThermoFisher, cat# 12-0459-42)

Staining was performed on ice for 30 minutes in FACS buffer, followed by two washes and resuspension in DAPI-containing FACS buffer. Fluorescence minus one (FMO) controls were used to define gates. Data were analyzed using FlowJo software (BD Biosciences), and populations were quantified based on consistent gating strategies across experiments.

### Cytokine Profiling

Conditioned media was collected from organoid co-cultures on day 7 of culture. For each sample, 20 μL of culture media was carefully harvested in triplicate and immediately stored at –20°C until further processing. Samples were submitted to the UCLA Immune Assessment Core for analysis using a 37-plex Luminex cytokine and chemokine panel.

All samples were run in technical triplicate, and cytokine concentrations were quantified against standard curves using the manufacturer’s software. Raw data were exported and analyzed using Microsoft Excel and R/RStudio. Cytokine values were normalized to the relevant control population and used to generate comparative heatmaps and bar plots. Samples falling below the detection threshold for a given cytokine were assigned a value of zero for downstream analysis.

### scRNASeq Cell Capture

Single-cell suspensions were generated from dissociated organoid co-cultures following FACS enrichment (GFP+ tumor or CD45+ immune cells). Cell suspensions were processed using the Chromium Controller and the Chromium Next GEM Single Cell 3’ v3.1 Reagent Kit (10x Genomics), following the manufacturer’s protocol. For each sample, up to 30,000 cells were targeted for capture. cDNA synthesis and library construction were performed according to 10x Genomics specifications. Libraries were sequenced on an Illumina NovaSeq 6000 platform at high depth to ensure saturation of the transcriptome complexity.

### scRNAseq Analysis

Raw sequencing reads were aligned and processed using the Cell Ranger pipeline (10x Genomics, v8.0.0) with default parameters. For human samples, reads were aligned to a standard GRCh38 reference genome obtained from 10x Genomics. Cell-by-gene UMI count matrices were generated for downstream analysis.

Initial data preprocessing was performed in R using the Seurat package (v5.0). Cells with fewer than 200 detected genes or greater than 10% mitochondrial gene content were excluded from the dataset. UMI counts were log-normalized and scaled with a size factor of 10,000. The top 2,000 most variable genes were identified, and principal component analysis (PCA) was applied to the scaled expression matrix. The number of principal components (PCs) used for downstream analysis was determined by identifying the maximum value between:

i. the squared standard deviation of each PC and
ii. the square root of (number of genes/number of cells + 1), squared.

Dimensional reduction and visualization were performed using UMAP based on the selected PCs. Cells were clustered using Seurat’s FindNeighbors and FindClusters functions with a resolution of 2.5.

### Cell Type Annotation

Cell type annotations were performed using Seurat’s reference-based mapping pipeline, following the recommended MapQuery protocol described in the Seurat documentation. Briefly, query datasets were normalized and integrated using SCTransform, and canonical correlation analysis (CCA) was used to identify anchors between the query and reference datasets. These anchors were then used to transfer cell type labels onto the query data.

For tumor compartments, cells were projected onto a previously published GBM single-cell meta-atlas (27). For immune compartments, reference-based mapping was performed using a previously published reference dataset, which includes matched gene expression and surface protein profiles across major immune lineages (26). Annotations were manually reviewed and refined using known marker gene expression if needed. Annotation results were used to define major cell types for all downstream analyses, including interaction modeling, gene set enrichment, and clonotype integration.

### GSEA

Gene set enrichment analysis was performed using the clusterProfiler package (v4.6.2) in R (59). Differential gene expression was computed using Seurat’s FindMarkers function with default Wilcoxon rank-sum testing. Resulting gene lists were ranked by average log fold change for use in GSEA. Pathway enrichment was assessed using the GSEA() function from clusterProfiler, querying against multiple gene set collections, including the MSigDB Hallmark, Reactome, and GO Biological Process databases. Gene sets with adjusted p-values < 0.05 were considered significantly enriched. Visualization of enriched pathways was performed using the dotplot function from the clusterProfiler suite.

### Ligand-Receptor Signaling Analysis

Cell-cell communication networks were inferred using the **CellChat** R package (v1.6.1), a tool for analyzing intercellular signaling based on known ligand-receptor interactions (60). Processed single-cell RNA-seq data were first subsetted by condition, and then normalized expression matrices and metadata were used to initialize CellChat objects.

CellChat’s built-in human ligand-receptor interaction database was used for all analyses. The workflow included identification of overexpressed genes and interactions, computation of communication probabilities, and aggregation of signaling networks. The computeCommunProb and computeCommunProbPathway functions were used to quantify global and pathway-specific communication, respectively. Sender-receiver relationships were visualized using circle plots, heatmaps, and hierarchical network diagrams. Differential pathway usage between conditions was computed using CellChat’s built-in statistical framework.

### TCR Sequencing and Analysis

T cell receptor (TCR) sequencing was performed using the 10x Genomics Chromium Single Cell V(D)J platform. Matched gene expression and TCR libraries were generated using the 10x Genomics Single Cell 5’ V(D)J kit according to the manufacturer’s instructions. Samples were processed in parallel for gene expression (GEX) and TCR enrichment using cellranger multi, which enables simultaneous demultiplexing, alignment, and quantification of both libraries. For V(D)J alignment, reads were mapped to a human TCR reference downloaded from the official 10x Genomics reference repository.

TCR contigs were assembled and annotated by Cell Ranger, including CDR3 sequences, clonotype IDs, and V(D)J gene usage. Productive, paired alpha-beta TCRs were retained for downstream analysis. Resulting filtered_contig_annotations.csv files were imported into R and integrated with single-cell gene expression data using the scRepertoire package (v1.7.1). TCR metadata (e.g., clonotype frequency, clonal expansion, and clone type) was added to Seurat objects for visualization and subgroup analysis. Clonotype dynamics, repertoire diversity, and overlap across samples or treatment conditions were assessed using scRepertoire and immunarch packages in R, adapting developer vignettes to build our analysis pipelines.

To investigate potential shared antigen recognition among clonotypes, GLIPH2 (Grouping of Lymphocyte Interactions by Paratope Hotspots) was used to cluster TCRs by sequence-based similarity (56). CDR3 sequences from productive, paired TCRs were input into GLIPH2, which outputs clusters of TCR CDR3 sequences predicted to recognize common epitopes. Output clusters were used to assess convergence within and across samples in our datasets.

## FIGURE LEGENDS: SUPPLEMENTARY FIGURES

**Supplementary Figure 1.**
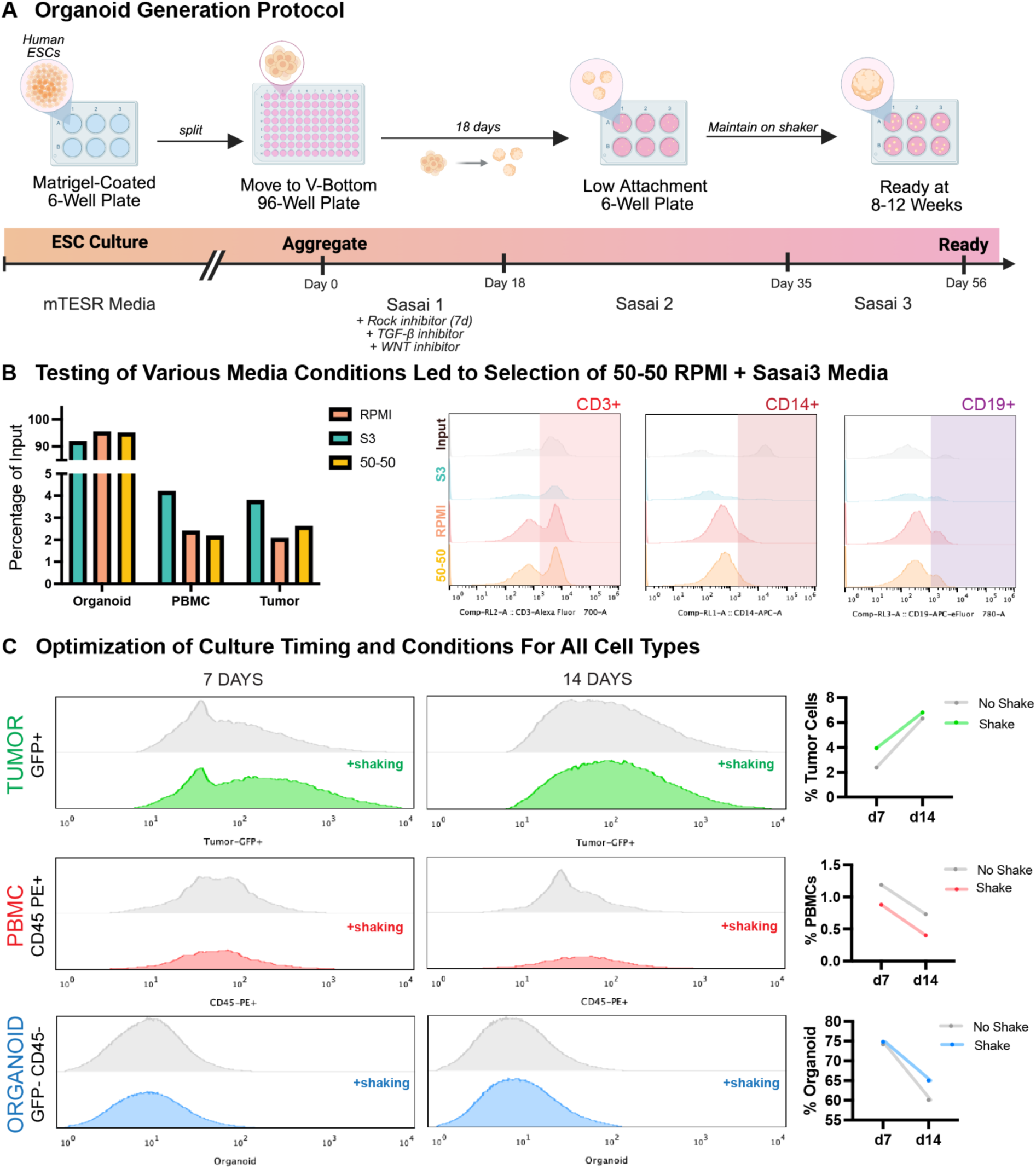
Organoid Generation and Optimization of iHOTT Culture Conditions A) Human embryonic stem cells (ESCs) were initially expanded in Matrigel-coated 6-well plates using mTESR media. Once confluent, cells were aggregated into V-bottom 96-well plates and cultured in Sasai1 media for 18 days. Organoids were then transferred to low-attachment 6-well plates and cultured in Sasai2 media until day 35, after which they were switched to Sasai3 media. Organoids were matured until day 56, at which point they were ready for tumor and PBMC co-transplantation. B) Three conditions—Sasai3, RPMI, and a 50:50 Sasai3/RPMI mixture—were tested for their ability to support the growth and viability of tumor cells, organoid tissue, and immune cell subtypes. (Left) The bar graph shows the percentage of cell recovery compared to the input of each compartment across media conditions. (Right) Flow cytometry assessing recovery of immune subpopulations across media conditions. The 50:50 Sasai3/RPMI condition was found to best support recovery of immune subpopulations and was used in subsequent experiments. C) Culture conditions and timing were assessed for iHOTT. Shaking improved the survival of tumor and organoid cells but did not benefit PBMC viability. Time-course analysis revealed that while tumor cells proliferated over time, PBMCs and organoid cells exhibited reduced viability at later time points. Day 7 was selected as the optimal endpoint for co-culture experiments to maximize cellular recovery across all compartments.

**Supplementary Figure 2:**
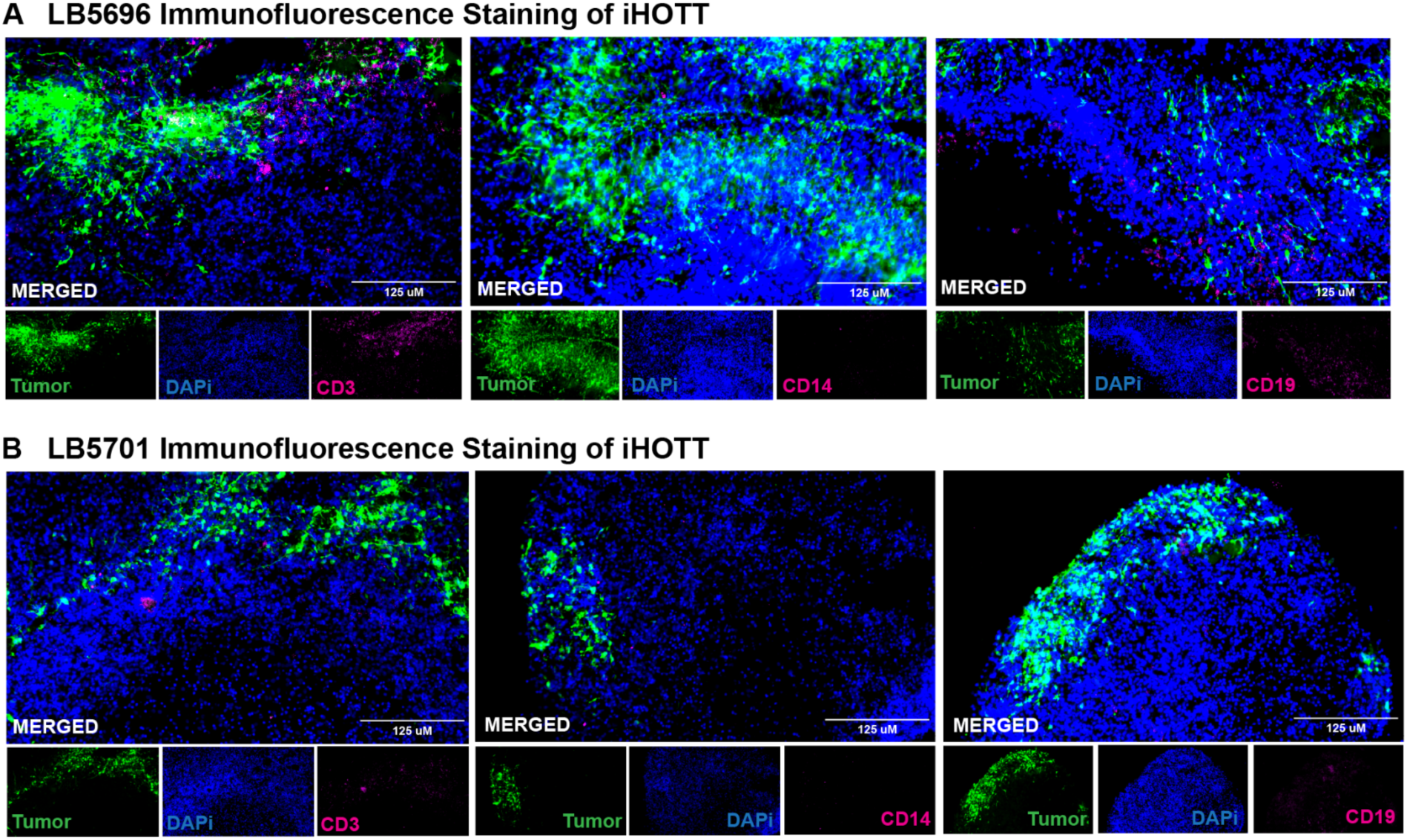
Immunofluorescence Staining of all Patients A) Immunofluorescence of LB5696 revealed infiltration of CD3+ T cells as the predominant immune population, with minor populations of CD14+ myeloid and CD19+ B cells. GFP+ tumor cells were observed invading into the organoid parenchyma. B) Immunofluorescence staining of iHOTT organoids from patient LB5701 showed a similar pattern, with robust CD3+ T cell infiltration, sparse CD14+ and CD19+ cells, and clear tumor integration into the organoid tissue.

**Supplementary Figure 3:**
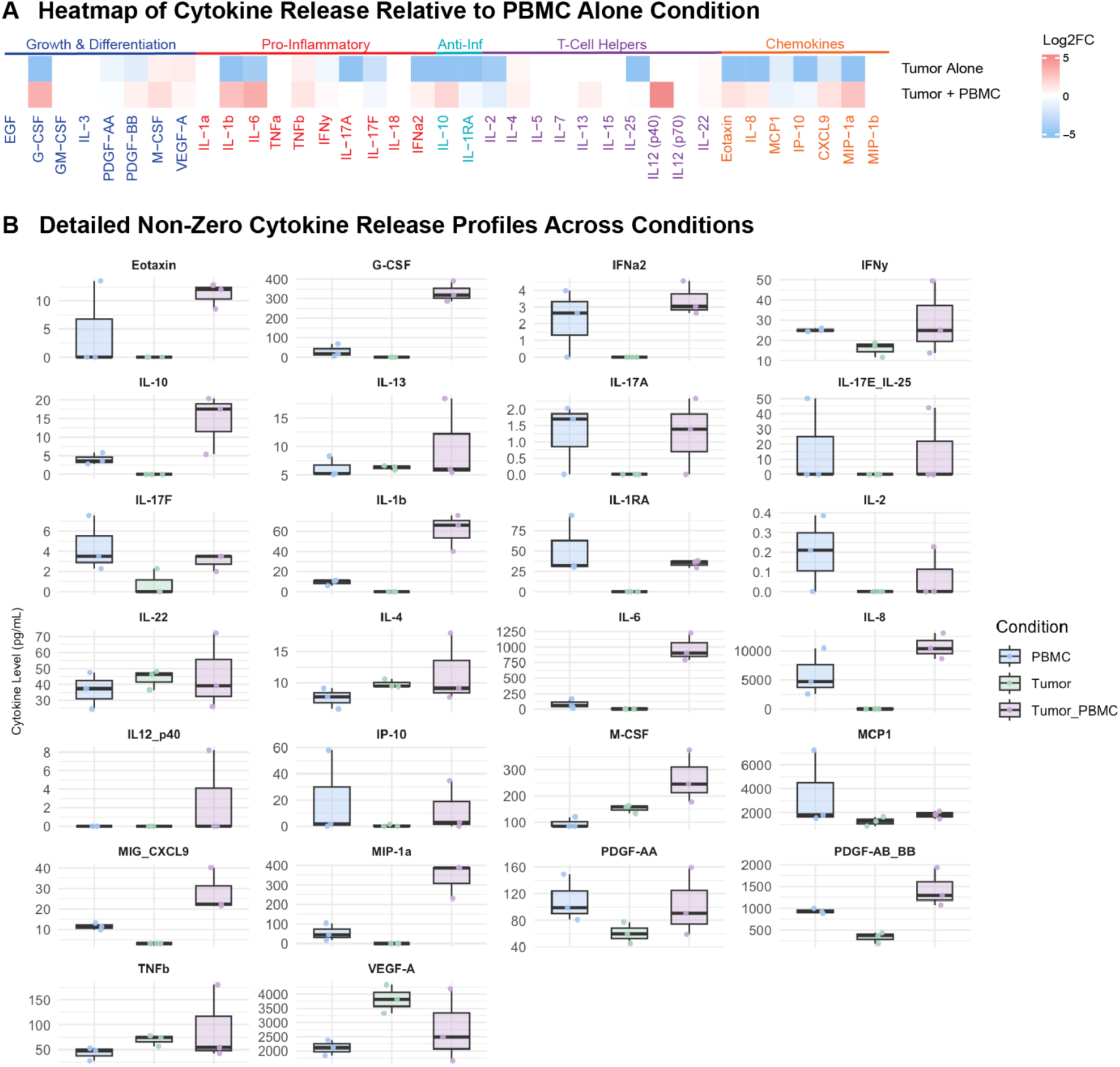
Cytokine Profiling Across Tumor-Only and PBMC-Only Controls Reflects the Importance of Co-Culture for Generating Cytokine Response A) Heatmap depicting cytokine expression profiles from LB5625 across three conditions: tumor-only, PBMC-only, and tumor + PBMC (iHOTT). Values are normalized to the PBMC-only condition. The tumor-only condition showed minimal cytokine secretion, while co-culture with PBMCs elicited a robust cytokine response, highlighting the necessity of immune-tumor interaction to recapitulate relevant cytokine signaling. B) Bar plots of individual cytokines detected in LB5625, limited to non-zero cytokines. Expression levels in tumor-only and tumor + PBMC conditions are shown relative to PBMC-only controls. Consistent with the heatmap, cytokine induction was substantially greater in the co-culture condition compared to the tumor alone, emphasizing the importance of the iHOTT system for modeling physiologic immune-tumor crosstalk.

**Supplementary Figure 4:**
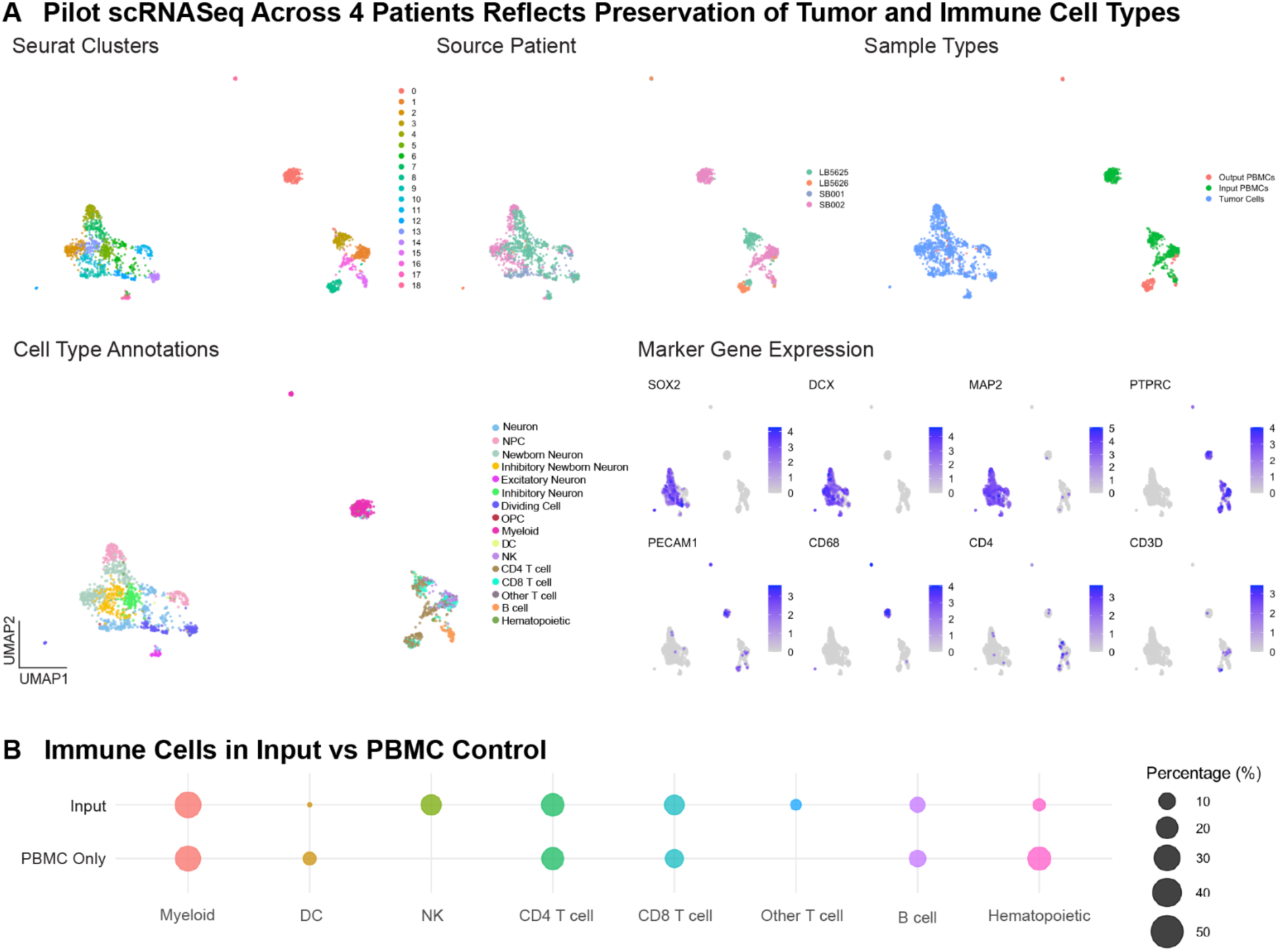
PBMC-Only Organoid Transplants Reflect Preservation of PBMCs in the Organoid System A) UMAPs of PBMC-only organoid co-cultures across four patient samples. Cells were annotated by Seurat clustering, patient origin, sample type, and cell type identity. Feature plots display canonical marker gene expression, confirming accurate immune cell type assignments. B) Comparison of immune cell composition across input PBMCs and PBMC-only co-cultures, showing maintenance of most immune cell types in the PBMC-only organoid condition.

**Supplementary Figure 5:**
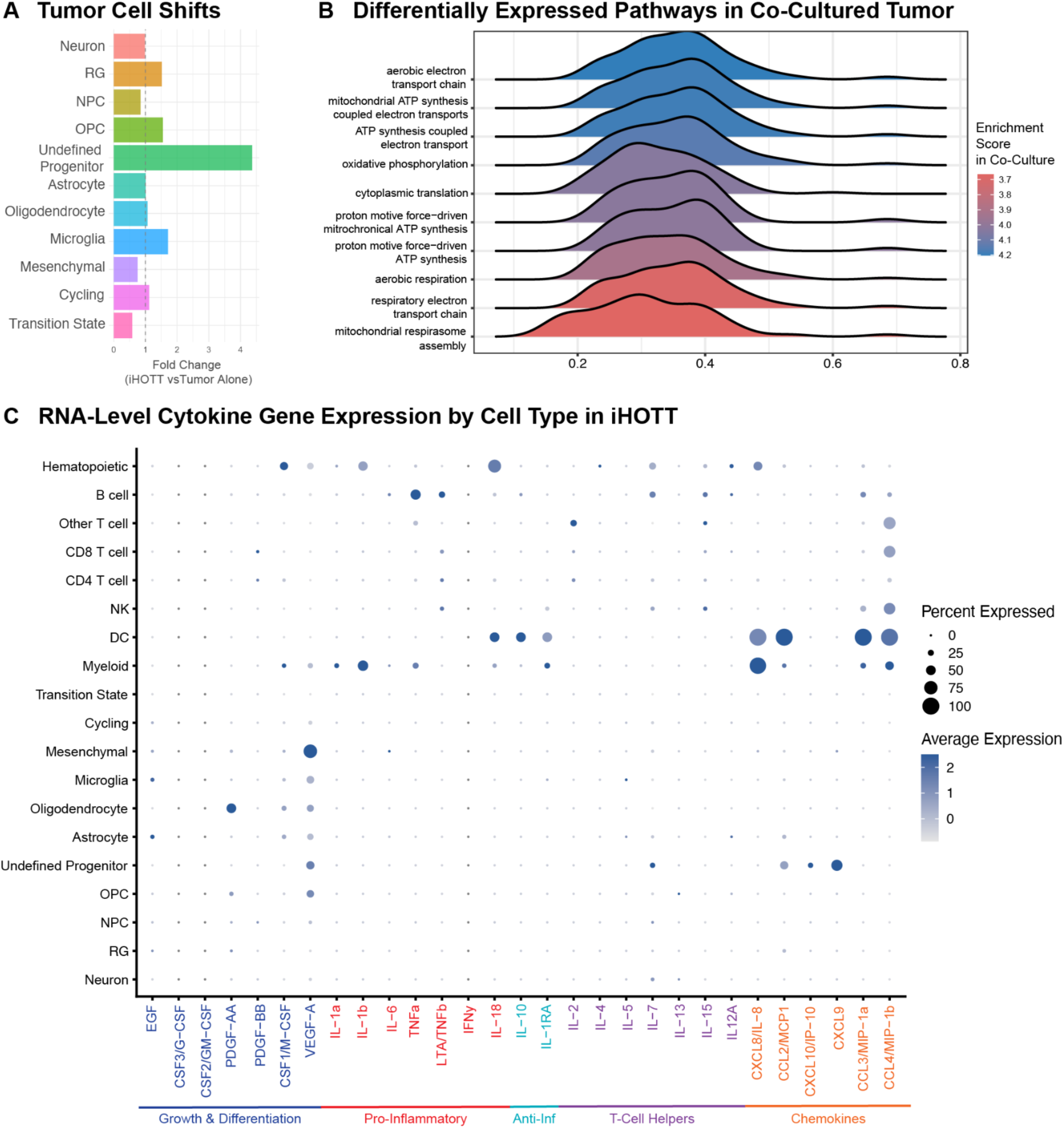
Examination of the Tumor Compartment Reveals Immune Co-Culture Induces Metabolic Changes in Tumor Cells A) Cell type distribution analysis comparing tumor cells from tumor-only organoid transplants and iHOTT co-cultures. B) Pathway analysis of differentially expressed genes in tumor cells revealed upregulation of multiple metabolic pathways in the iHOTT condition. These findings highlight the importance of studying tumor cells in the context of the immune microenvironment to capture relevant metabolic adaptations. C) Gene expression of available cytokines in our single-cell RNAseq dataset for iHOTT is shown, stratified by cell type.

**Supplementary Figure 6:**
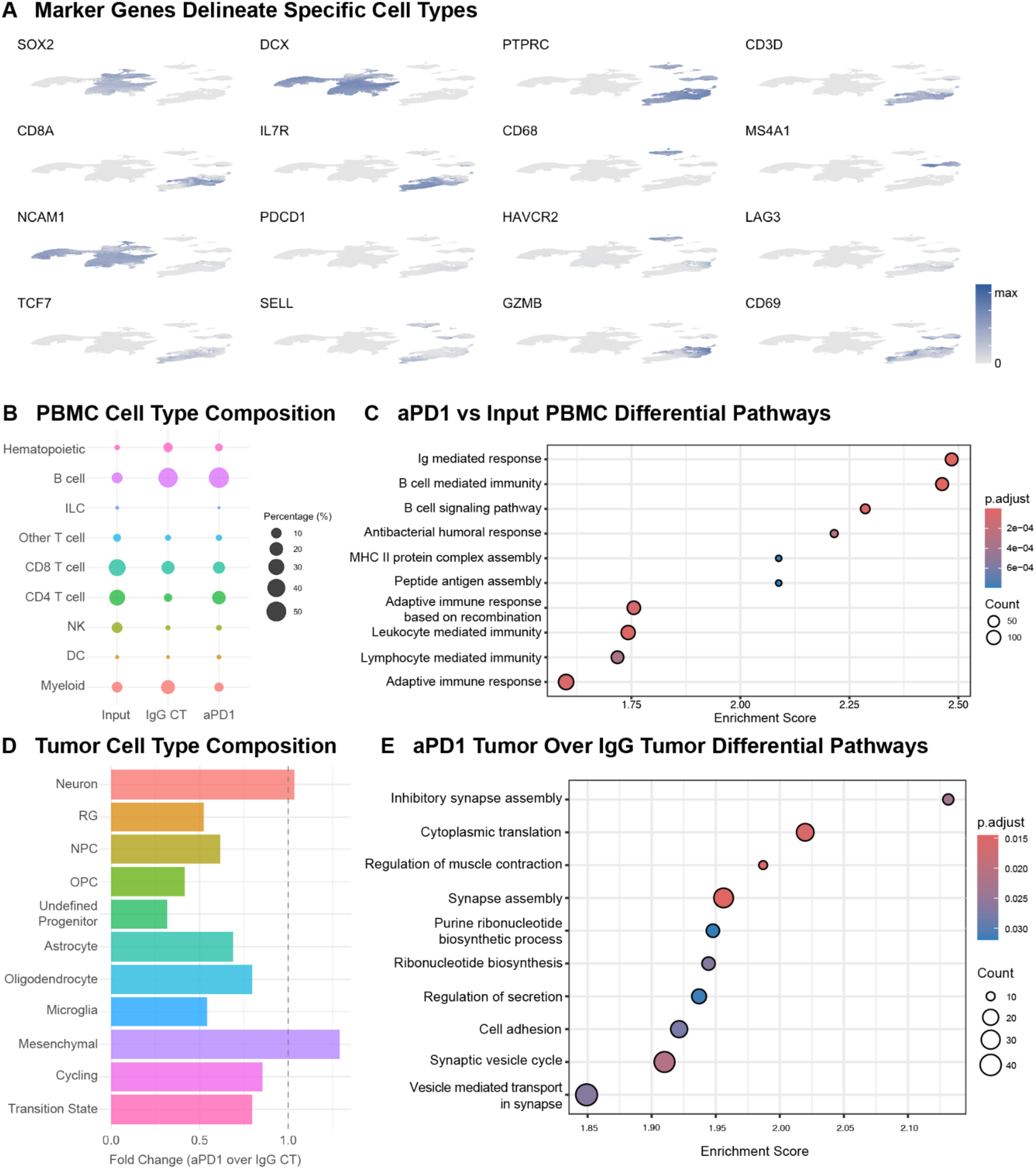
Further Analysis of Pembrolizumab Treatment Effects on iHOTT Tumor and Immune Compartments A) Expression of marker genes used to annotate tumor and immune clusters, defining cell types annotated in Figure 3. B) PBMC composition across input, IgG control, and pembrolizumab-treated iHOTT samples reflects that all major immune populations were maintained, and that there was subtle expansion of B and T cell populations. C) GSEA comparing pembrolizumab-treated PBMCs to input PBMCs revealed enrichment of immune activation pathways, consistent with a tumor-driven, treatment-responsive immune state. D) Tumor cell type composition is shown across treatment conditions E) GSEA of tumor cells revealed increased expression of synaptic, transport, and biosynthesis pathways following pembrolizumab treatment. These findings suggest that tumor cells exhibit a reactive transcriptional response to immune engagement in the iHOTT system.

**Supplementary Figure 7:**
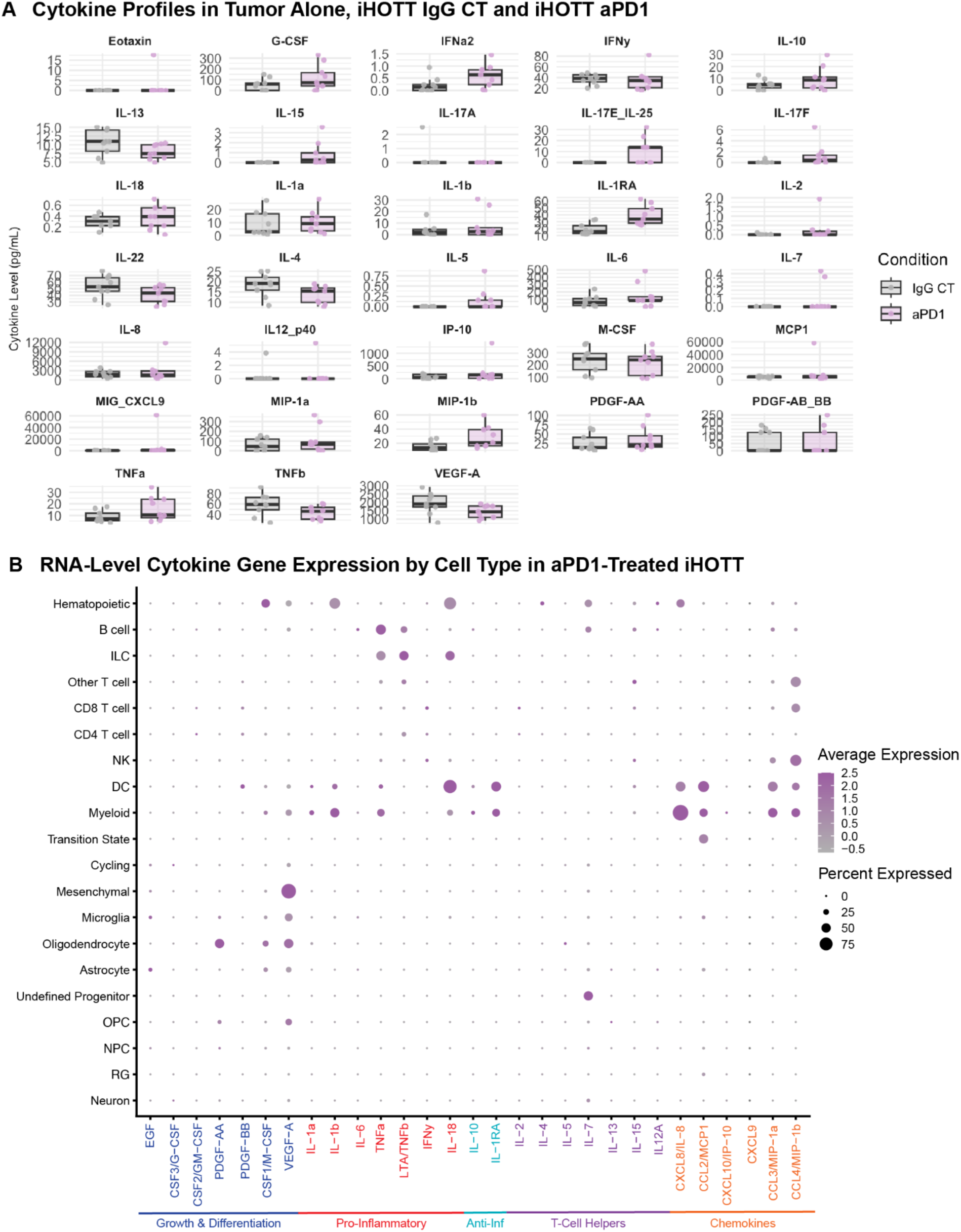
Detailed Cytokine Release Profiles Reveal Enhanced Polarization in Pembrolizumab-Treated Conditions A) Bar plots depicting cytokine secretion levels across IgG control and pembrolizumab-treated iHOTT samples for all cytokines measured. Across multiple tumors, pembrolizumab treatment resulted in further polarization of the cytokine response trend seen in the control condition. These profiles underscore the immune-activating effects of PD-1 blockade within the iHOTT co-culture system. B) Single-cell RNA-seq data for matched available cytokines after pembrolizumab treatment is shown, stratified by cell type.

**Supplementary Figure 8:**
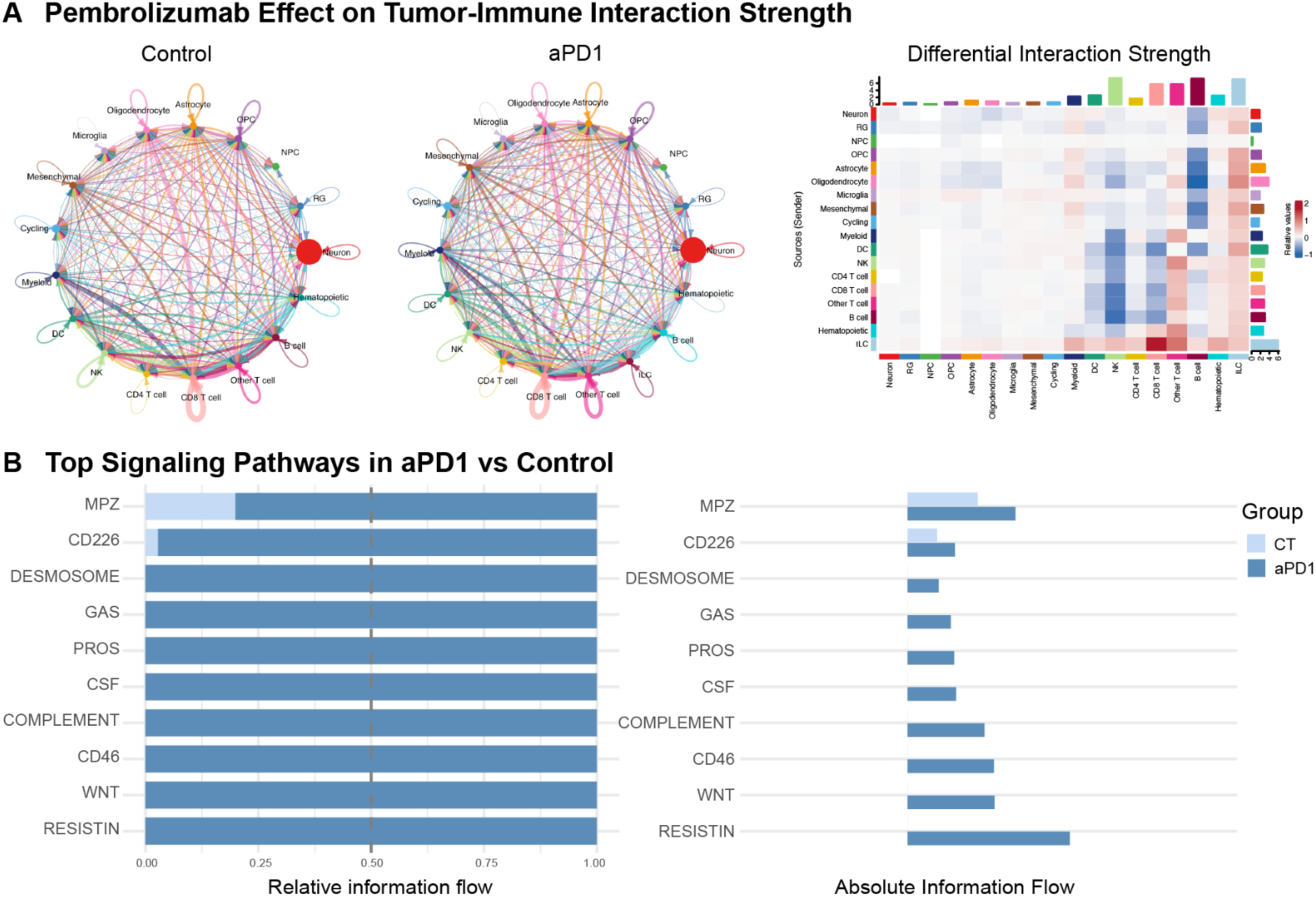
Signaling Interactions Between Tumor Cells and PBMCs Reflect Immune Signaling Upregulation in Pembrolizumab-Treated iHOTT Compared to Control A) (Left) CellChat interaction plots comparing global ligand-receptor signaling networks between IgG control and pembrolizumab-treated iHOTT samples. (Right) Differential interaction analysis revealed upregulated interactions between CD8 T cells and ILCs, other T cells, and all immune cell types, and ILCs and all cell types in pembrolizumab-treated samples. B) Top signaling pathways enriched in pembrolizumab-treated iHOTT included immune-related pathways such as RESISTIN, WNT, CD45, complement, and others, highlighting the activation of diverse immune communication programs following PD-1 blockade.

**Supplementary Figure 9:**
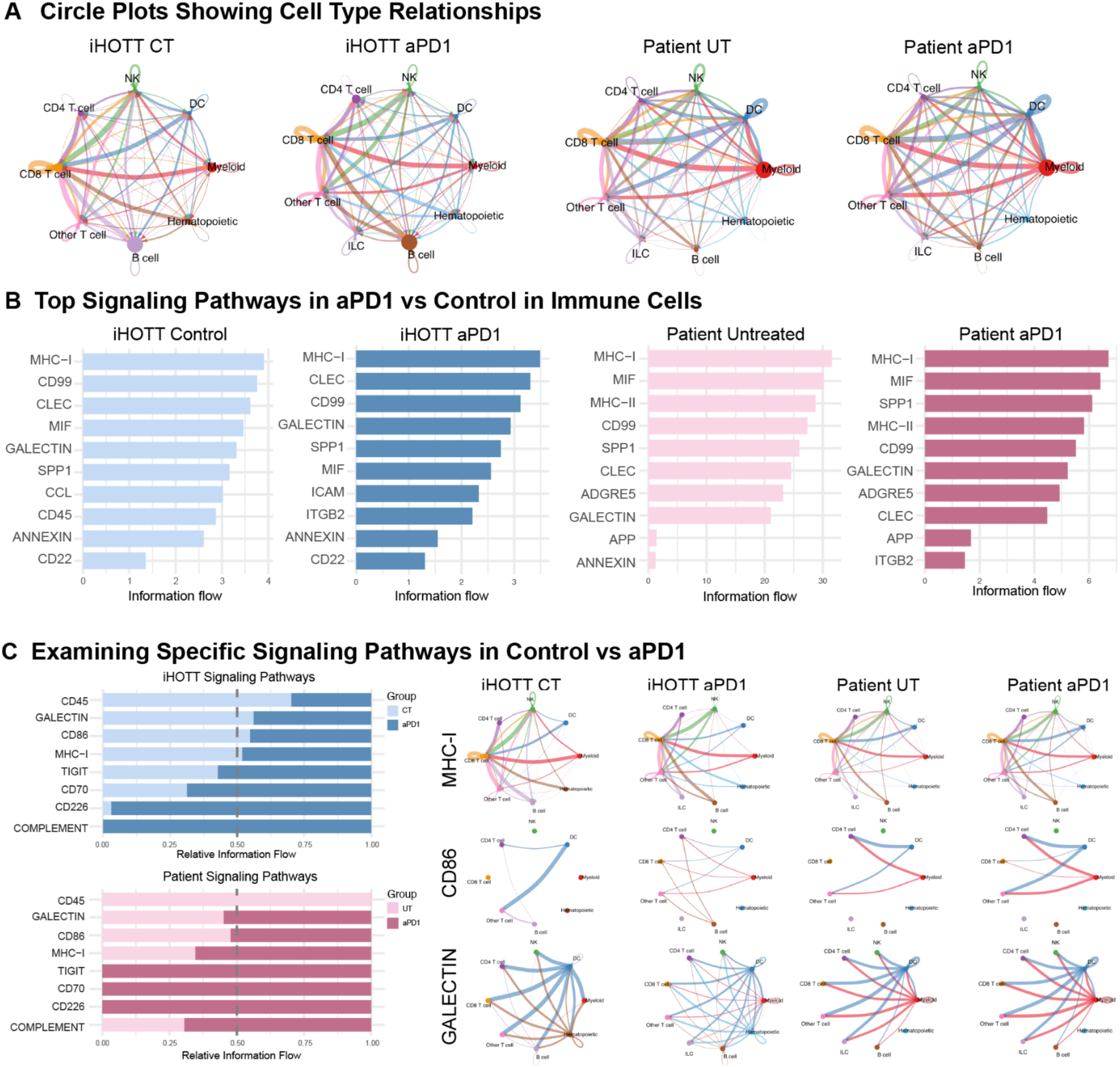
Comparative Analysis of PBMC Signaling Interactions in iHOTT and Pembrolizumab-Treated Patient Samples A) CellChat circle plots comparing ligand-receptor interactions in PBMC compartments from iHOTT and patient tumor samples. Both systems show increased interactions involving ILCs and other T cells following pembrolizumab treatment, highlighting consistent immune activation across models. B) Analysis of signaling pathways revealed MHC-I as a core pathway active across all conditions. Additional immune pathways were also activated across all conditions, as shown. C) Examination of immune checkpoint–associated signaling showed concordant trends in iHOTT and patient samples. CD45 signaling was predominantly active in IgG/untreated conditions, while Complement and CD226 signaling were enriched in pembrolizumab-treated conditions. Similar immune cell types were involved in mediating these pathways in both systems, emphasizing strong concordance between iHOTT and the human treatment context.

**Supplementary Figure 10:**
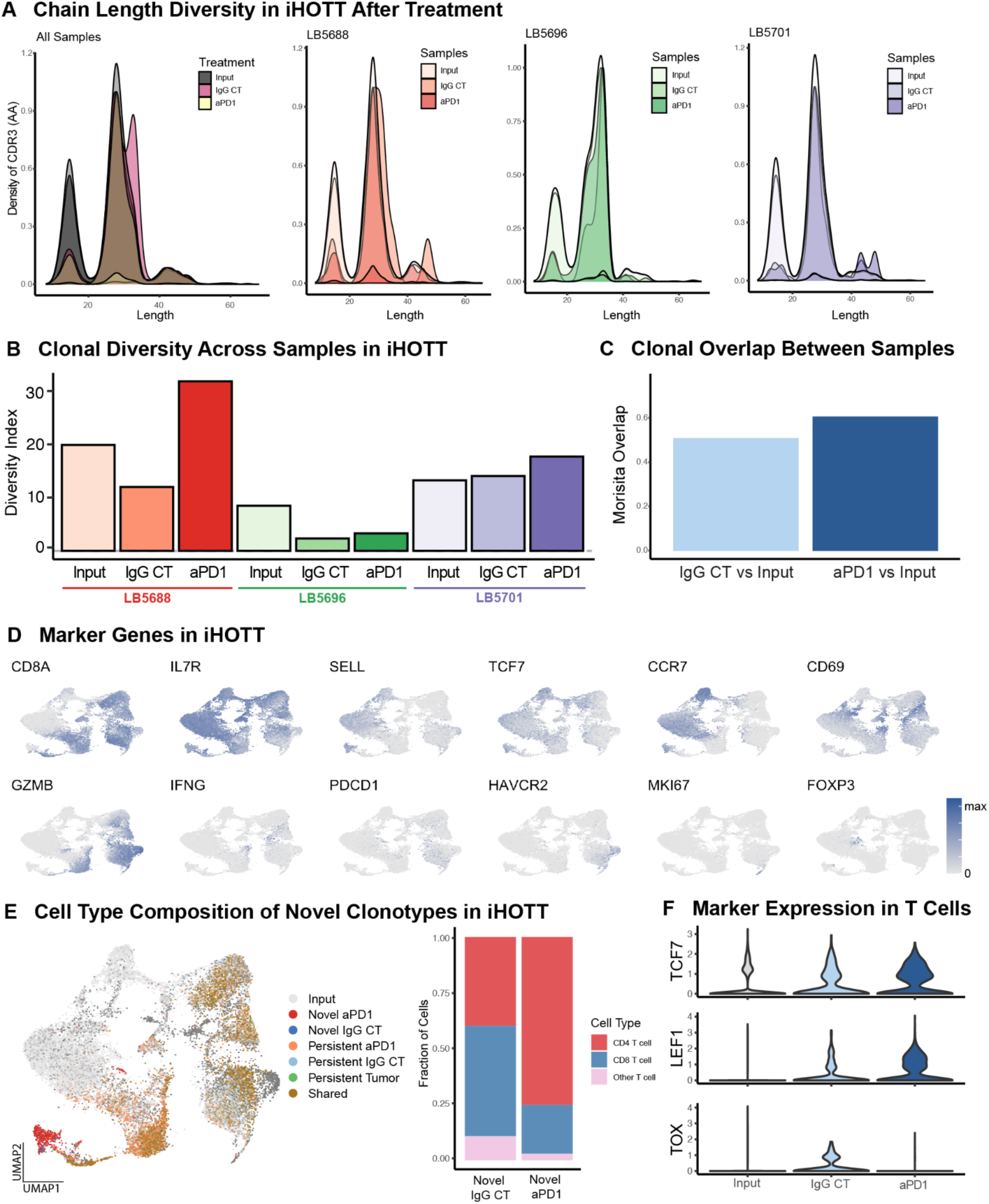
Detailed Clonotype Analysis in iHOTT Reveals Diversity and Transcriptional Signatures of Novel T Cell Clones A) Chain length density plots from TCR sequencing of iHOTT samples show shifts in chain length distribution following pembrolizumab treatment, indicating possible clonal remodeling of the T cell repertoire. B) Clonal diversity across individual iHOTT samples shows that pembrolizumab-treated samples exhibited increased TCR diversity relative to input and IgG control. C) Clonal overlap analysis revealed a comparable proportion of persistent clonotypes in pembrolizumab-treated samples relative to control. D) Marker gene expression is shown projected on our T cell UMAP in iHOTT< reflecting markers of activation, exhaustion, and cell types. E) Left: UMAP of reclustered T cells colored by clonotype persistence across treatments **within the same patient.** Input refers to clonotypes seen only in the input sample for that patient. Novel aPD1 or CT refers to clonotypes that were novel in aPD1 or control for that patient. Persistent aPD1 or CT refers to clonotypes that were persistent across input and either aPD1 or CT respectively. Persistent tumor reflects clonotypes seen in both CT and aPD1 for a given patient but not seen in the input. Finally, shared refers to clonotypes seen in all 3 conditions for a given patient. Right: Cell type composition of novel clonotypes showed CD4+ T cells as the dominant source in both conditions, with a pronounced enrichment in pembrolizumab-treated samples. F) Violin plots of TCF7, LEF1, and TOX expression among T cells revealed increased expression of progenitor T cell precursor markers (TCF7, LEF1) and reduced expression of exhaustion marker TOX in pembrolizumab-treated samples, consistent with a reinvigorated T cell state.

**Supplementary Figure 11:**
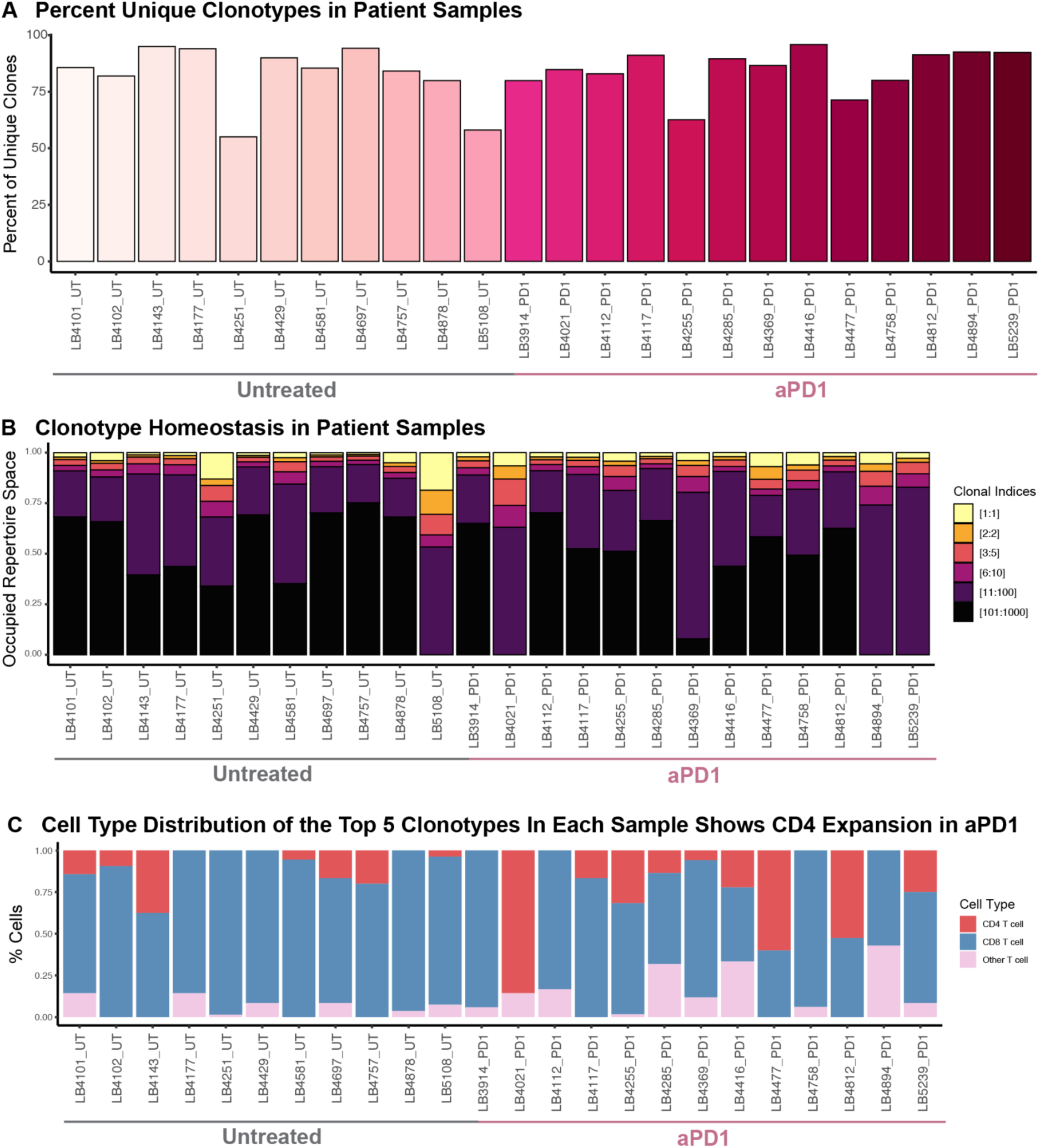
Patient-Level Clonotype Repertoire Analysis Highlights CD4 T Cell Enrichment in Pembrolizumab-Treated Tumors A) Quantification of the percentage of unique TCR clonotypes across patient tumor samples, comparing untreated and pembrolizumab-treated cohorts. B) Clonotype homeostasis analysis revealed significant heterogeneity in repertoire structure across patients. C) Sample-level analysis of the top 5 clonotypes in each patient tumor showed that dominant clones in pembrolizumab-treated samples were more frequently composed of CD4+ T cells and Other T cell populations, suggesting treatment-associated reshaping of the T cell repertoire toward these subsets.

